# Intranasal delivery of NS1-deleted influenza virus vectored COVID-19 vaccine restrains the SARS-CoV-2 inflammatory response

**DOI:** 10.1101/2022.10.03.510566

**Authors:** Liang Zhang, Yao Jiang, Jinhang He, Junyu Chen, Ruoyao Qi, Lunzhi Yuan, Tiange Shao, Congjie Chen, Yaode Chen, Xijing Wang, Xing Lei, Qingxiang Gao, Chunlan Zhuang, Ming Zhou, Jian Ma, Wei Liu, Man Yang, Rao Fu, Yangtao Wu, Feng Chen, Hualong Xiong, Meifeng Nie, Yiyi Chen, Kun Wu, Mujing Fang, Yingbin Wang, Zizheng Zheng, Shoujie Huang, Shengxiang Ge, Shih Chin Cheng, Huachen Zhu, Tong Cheng, Quan Yuan, Ting Wu, Jun Zhang, Yixin Chen, Tianying Zhang, Hai Qi, Yi Guan, Ningshao Xia

**Affiliations:** State Key Laboratory of Molecular Vaccinology and Molecular Diagnostics; National Institute of Diagnostics and Vaccine Development in Infectious Diseases, School of Public Health & School of Life Sciences, Xiamen University, Xiamen 361102, Fujian, China; Tsinghua-Peking Center for Life Sciences, Laboratory of Dynamic Immunobiology, School of Medicine, Tsinghua University, Beijing, 100084, China; State Key Laboratory of Cellular Stress Biology, School of Life Sciences, Xiamen University, Xiamen, 361102, Fujian, China; Xiang An Biomedicine Laboratory, Xiamen 361102, Fujian, China; State Key Laboratory of Emerging Infectious Diseases, School of Public Health, Li Ka Shing Faculty of Medicine, The University of Hong Kong, Hong Kong 999077, China; Guangdong-Hong Kong Joint Laboratory of Emerging Infectious Diseases/Joint Laboratory for International Collaboration in Virology and Emerging Infectious Diseases, Joint Institute of Virology (STU/HKU), Shantou University, Shantou 515063, China

**Author notes:** These authors contributed equally to this work (Liang Zhang, Yao Jiang, Jinhang He, Junyu Chen, Ruoyao Qi, Lunzhi Yuan, Tiange Shao). Corresponding authors. Correspondence (N. Xia), (Y Guan), (H Qi), (T. Zhang), (Y. Chen), (J Zhang), (T Wu).

**Keywords:** SARS-CoV-2, COVID-19 vaccine, live attenuated influenza virus vector, NS1-deleted, intranasal vaccine, innate immunity, trained immunity, tissue-resident memory T cells, broad-spectrum, respiratory mucosal immunity

## Abstract

The emergence of SARS-CoV-2 (Severe Acute Respiratory Syndrome Coronavirus-2) variants and “anatomical escape” characteristics threaten the effectiveness of current coronavirus disease (COVID-19) vaccines. There is an urgent need to understand the immunological mechanism of broad-spectrum respiratory tract protection to guide broader vaccines development. In this study, we investigated immune responses induced by an NS1-deleted influenza virus vectored intranasal COVID-19 vaccine (dNS1-RBD) which provides broad-spectrum protection against SARS-CoV-2 variants. Intranasal delivery of dNS1-RBD induced innate immunity, trained immunity and tissue-resident memory T cells covering the upper and lower respiratory tract. It restrained the inflammatory response by suppressing early phase viral load post SARS-CoV-2 challenge and attenuating pro-inflammatory cytokine (*IL-6, IL-1B*, and *IFN-*γ) levels, thereby reducing excess immune-induced tissue injury compared with the control group. By inducing local cellular immunity and trained immunity, intranasal delivery of NS1-deleted influenza virus vectored vaccine represents a broad-spectrum COVID-19 vaccine strategy to reduce disease burden.

## Introduction

SARS-CoV-2 (severe acute respiratory syndrome–related coronavirus) has infected over 580 million people, claimed over 6 million lives, and caused a dramatic loss to human society as of July 2022. SARS-CoV-2 invades the host by binding to angiotensin converting enzyme 2 (ACE2), a high-affinity receptor on respiratory epithelial cell surfaces. Currently, 38 vaccines have been approved for use, most of which are vaccinated by intramuscular injection. The protective effects are mainly derived from the neutralizing antibody targeting spike antigen, and large-scale vaccination has effectively reduced SARS-CoV-2 symptomatic infection, hospitalization and death ^1–3^. Antibody levels in the respiratory tract are 200-500 times lower than that in circulation ^4^ which leads to “anatomical escape” of SARS-CoV-2 in the upper respiratory tract since it is difficult to completely block the infection, especially after the peak vaccine-induced immune response period ^5–7^. Furthermore, escape variants are emerging in an endless stream; for example, the Omicron mutant strain has the most significant changes in antigenicity and the immunity conferred after vaccinations and natural infection ^8,9^. Finally, the virus is also hosted by several animal reservoirs such as minks, cats, deer’s ^10,11^. These realities portend that coronavirus disease (COVID-19) will coexist with humans for many years and will pose a continuing threat. Therefore, the development of broad-spectrum COVID-19 vaccines that rely on various immune mechanisms and different technical routes should be encouraged.

Theoretically, local protective immune factors in the respiratory tract should respond to SARS-CoV-2 infection in a timelier manner than effectors present in the peripheral lymph nodes and blood. Therefore, the development of COVID-19 vaccines via respiratory inoculation has become a hot pipeline shown positive effects in preclinical animal experiments, including vaccines based on adenovirus vectors, vesicular stomatitis virus vectors and other viral vectors ^12–14^. An Ad-vectored trivalent COVID-19 vaccine expressing spike-1, nucleocapsid, and RNA-dependent RNA polymerase (RdRP) antigens from Zhou Xing et al. shows a good broad-spectrum protective effect against a variety of SARS-CoV-2 variant strains by intranasal vaccination ^15^. In addition, trained immunity not dependent on specific antigen epitope also plays an important role in the broad-spectrum efficacy of this respiratory mucosal vaccine ^15^. It would be difficult to achieve sterilizing immunity against SARS-CoV-2 infection by vaccination as the emerging variants and “anatomical escape” characteristics. Instead, inducing a local immune regulation mechanism that preventing an excessive inflammatory response in the respiratory tract will achieve broad-spectrum protection and reduce the COVID-19 disease burden. The broad-spectrum protective effect of NS1-impaired influenza virus on heterologous influenza virus challenge is independent of virus clearance and deserves attention ^16^. Intranasal immunization with NS1-truncated virus (A/PR8/NS124) induces stronger effector T-cells and certain immunoregulatory mechanisms compared with wild-type H1N1 influenza strain (A/PR8/NSfull), which protects the organism against lethal heterologous A/Aichi/2/68 (H3N2) influenza virus challenge through significant attenuation of inflammation and pathology without inhibiting viral load ^16^. These studies suggest raise a novel/special hypothesis that the vaccine-induced protective effect may be derived from immune regulation in the respiratory tract to prevent excessive inflammation, but not limited to block viral infection or suppress viral levels.

Modification of NS1 protein is a promising approach for the development of live-attenuated influenza viral vectors ^17^. We previous developed an intranasal spray vaccine based on the NS1-deleted H1N1 vector carrying the gene encoding SARS-CoV-2-RBD (dNS1-RBD) ^18,19^, which is currently undergoing Phase III clinical trials in several countries [ChiCTR2100051391]. This vaccine prevents COVID-19 induced by Prototype, Beta and Omicron of SARS-CoV-2 challenge in hamster models in the absence of detectable neutralizing antibodies ^18^. The immunological mechanisms that provide broad-spectrum protection remains unclear.

In this study, we investigated the protective immune response induced by dNS1-RBD. The broad-spectrum protective immunity induced by this vaccine mainly includes the following aspects: i), innate immunity, various cytokines and chemokines containing antiviral functions or initiating local immune responses were detected in lung tissue within 24 hours after vaccination; alveolar macrophages (AMs), dendritic cells (DCs), and NK cells (NK) were also activated; ii), trained immunity that realizes the memory response of innate immunity by reprogramming the chromatin accessibility landscape which reshapes the immune response profile upon SARS-CoV-2 infection, with attenuation of pro-inflammatory factors and pathways; iii), striking local T cell responses covering the upper and lower respiratory tract. Tissue resident T cells were detected in the nasal-associated lymphoid tissue (NALT) and the lung which supports the long-term protective effects. This intranasal vaccine represents an effective broad-spectrum COVID-19 vaccine strategy by inducing specific and non-specific protective immunity, particularly in the respiratory tract.

## Results

### dNS1-RBD immunization systemically activated antiviral innate immune pathways in the lung

The innate immune response elicited by dNS1-RBD was explored by collecting 15 samples of mouse lung tissue for RNA-seq analysis, including 12 samples from 1-, 7-, 14- and 28 days post-immunization (d.p.im), and three samples without treatment as the control group (**Fig 1A**). Overall, gene expression levels were clustered in principal component analysis (PCA) space along the PC1 and PC2 coordinates (**Fig 1B**). Many genes from the innate immune response and cytokine-related pathways were rapidly upregulated on the first day after immunization and gradually returned to a resting state over time (**Fig 1C**). Differentially expressed genes (DEGs) extracted from the dNS1-RBD group *vs*. the control group were taken as the key factors and displayed in a heatmap; significant transcriptional changes were observed in the lung during the early phase (Day 1- and Day 7 group) after vaccination and the effect gradually decreased in the late phase (Day 14- and Day 28 group) (**Fig 1D**, left). These genes were mostly activated through dynamic curves that peaked from 1 d.p.im to 7 d.p.im which were further divided into four clusters (**Fig 1D**, middle). The genes from the Cluster 1 to 4 were highly enriched in innate immune response-related pathways such as cytokine production, chemokine signaling pathway, Nod-like receptor signaling pathway and TNF signaling pathway based on Gene Ontology (GO) and Kyoto Encyclopedia of Genes and Genomes (KEGG) enrichment analyses (**Fig 1D**, right). Significant elevation of interferon-related genes (*Ifit2, Ifna1, Ifna2, Ifnb1*) and transcription factor *Stat1* was detected at 1 d.p.im and returned to the resting state at 7 d.p.im (**Fig 1D**, right). Our previous study showed that several lung tissue cytokines (*interferon alpha, IFN-*α *and Interferon gamma, IFN-*γ) are rapidly upregulated by dNS1-RBD at 1 d.p.im and are significantly higher than those in wild-type H1N1/CA04 infected mice ^18^. These results may be associated with NS1 deletion, which acts as an antagonist of host type-I interferon responses. These results suggest that dNS1-RBD systemically activates antiviral innate immune responses in mice lungs which may support the rapid establishment of protective immunity after vaccination.

**Fig 1.**
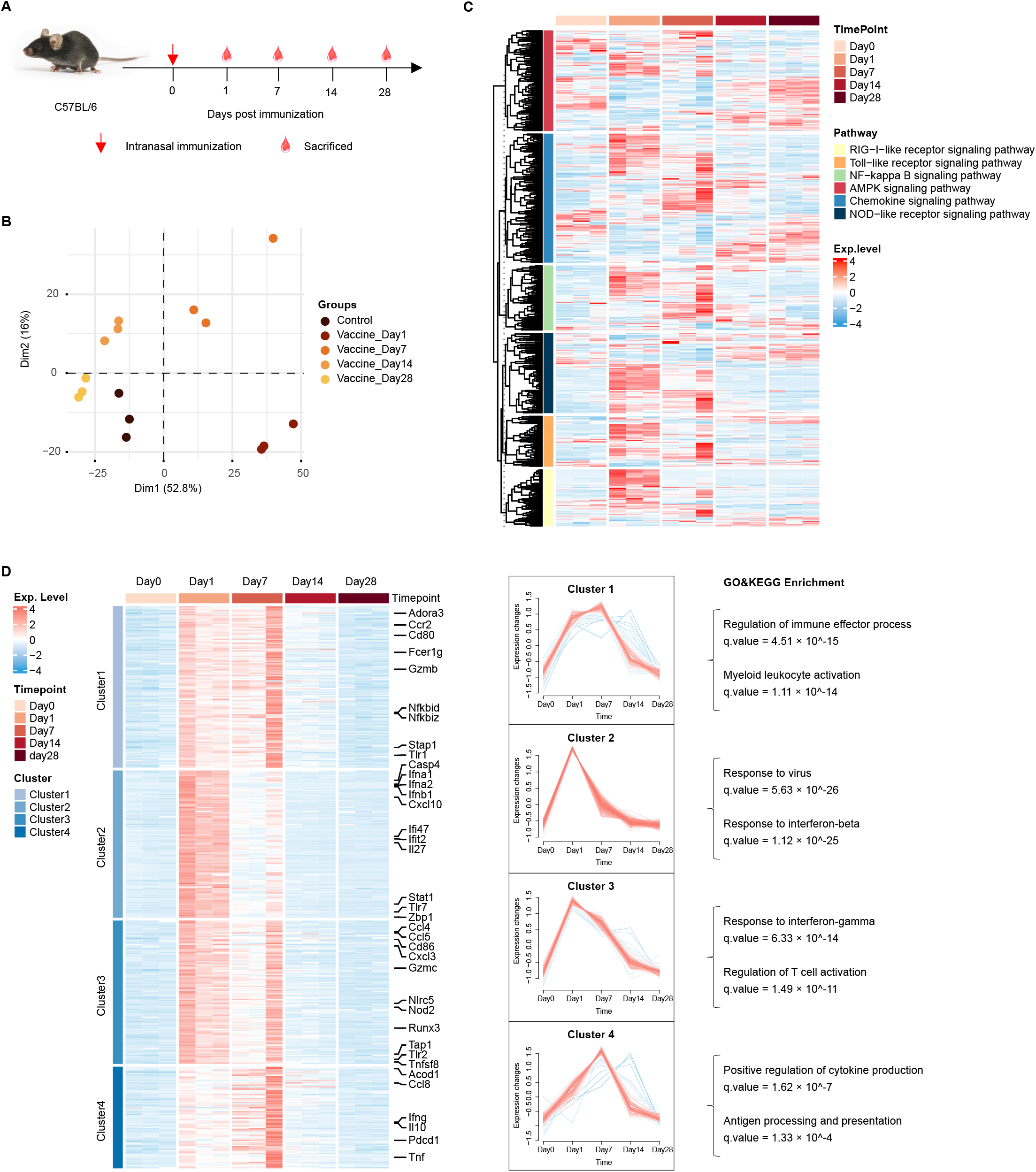
Transcriptional dynamics show activated antiviral innate immune responses in the lung induced by dNS1-RBD. (A) Experimental Schema. (B) Principal component analysis (PCA) of the data collected from the lung of C57BL/6 mice at 1-, 7-, 14-, and 28-days post-immunization. n = 3 mice/group. (C) Heatmap showing KEGG pathway-based enrichment analysis of DEGs. (D) Heatmap showing the dynamic expression patterns of DEGs. Expression trends of the genes in the four clusters are shown in the right part.

### dNS1-RBD immunization established a local innate immune barrier by activating alveolar macrophages, DCs, NK cells, and virtual memory T cells (T_VM_)

Immunization by dNS1-RBD and the empty vector (dNS1-Vector) control provides a protective effect in the early stage (1 d.p.im) ^18^. The local innate immune responses were characterized in the lungs at the cellular level by intranasal immunization of C57BL/6 mice with a single dose of dNS1-RBD or dNS1-Vector. Lung immune cells were harvested and analyzed at different time points by flow cytometry (**Fig S1A, B**). Alveolar macrophages are the first-line immune cells in lung tissue exposed to pathogens, are critical for the early control of SARS-CoV-2 and are considered to be the driver of cytokine storms ^20^. After prime immunization, the number of AMs slightly decreased, but recovered on day 5 (**Fig 2A**). Major histocompatibility-II (MHC-II) levels rapidly increased in the AMs persisted for over 5 days, which was comparable between dNS1-Vector group and dNS1-RBD group (**Fig 2B**). The CD11b^high^ population significantly increased in AMs at 3 d.p.im (P < 0.001) which may be related to the properties of H1N1 vector (**Fig 2C-E**) ^21,22^. CD11b expression on the macrophage surface is upregulated after influenza A virus infection which promotes macrophage migration to the niche of infection and helps reduce the inflammatory pathological response ^21,22^. Plasmacytoid dendritic cells (pDCs) also played an important role in the early antiviral process by producing large amounts of antiviral cytokines; they were activated at 1 d.p.im and lasted for several days (**Fig 2F**). IFN-γ-secreting NK cells were observed in the dNS1-RBD group after 3 d.p.im, whereas they were observed at 5 d.p.im in the dNS1-Vector group (**Fig 2G, H**). As expected, the IFN-γ response was 20-fold higher in percentage in the lungs than that in the spleen (P < 0.01) (**Fig S2C**).

**Fig 2.**
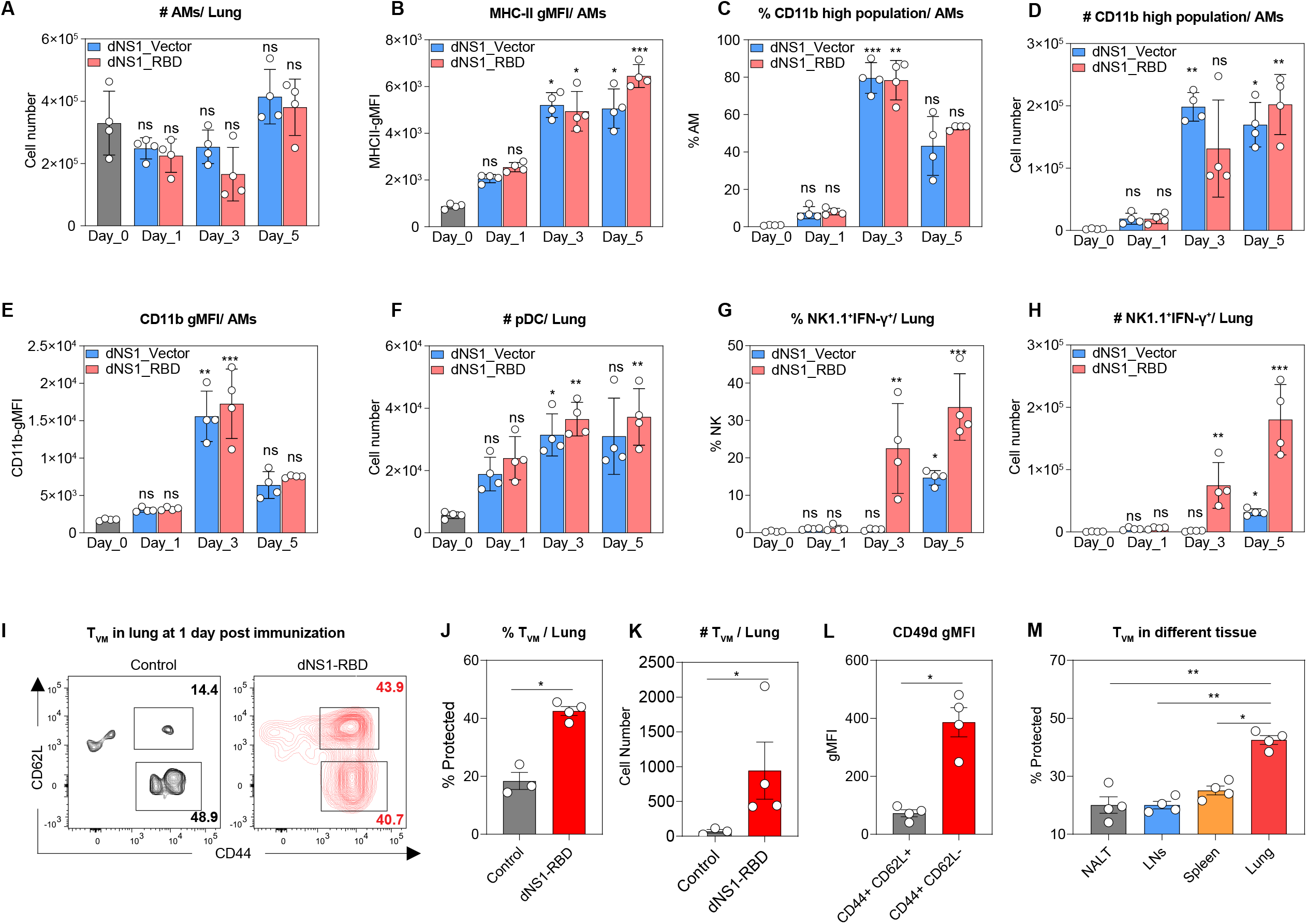
dNS1-RBD activates alveolar macrophages, DCs, NK cells, and virtual memory T Cells in the Lung. (A) Absolute number of AMs in the lung. (B) gMFI of MHC-II expression on AMs. (C-E) Frequency (C), absolute cell number (D), and gMFI (E) of CD11b^hi^ AMs in the lung. (F) Absolute number of pDCs in the lung. (G-H) Frequency (G) and absolute cell number (H) of IFN-γ^+^ NK1.1^+^ cells in the lung. (I) Representative flow cytometry contour plots for T_VMs_ in the lung. (J-L) Frequency (J), absolute cell number (K), CD49d expression expressed as gMFI (L) of T_VMs._ (M) Frequency of T_VMs_ in different tissues (NALT/LNs/spleen/lung). Data are presented as mean ± SEM. Statistical analysis for (A, B, C, D, E, F, G, and H) were Kruskal-Wallis tests with Dunn’s multiple comparisons test. Statistical analysis for (J, K, L, and M) were Mann-Whitney tests. ns, non-significant. *p < 0.05, **p < 0.01, ***p < 0.001. n = 3–4 mice/group. AMs: alveolar macrophages; gMFI: Geometry Mean Fluorescence Intensity; MHC-II: major histocompatibility-II; pDCs: plasmacytoid dendritic cells; NK cells: natural killer cells; IFN-γ: interferon-gamma; T_VMs_: Virtual memory T cells. NALT: Nasal-associated lymphoid tissue; LNs: Lymph nodes.

Previous reports indicated that virtual memory (VM) cells are capable of mediating both antigen-specific and bystander protective immunity against infection ^23^. Memory CD8^+^ T cells of VM phenotype have been reported to play a major role in the response of aged mice to lung infections ^24^. CD8^+^ T cell infiltration in different tissues was detected at 1 d.p.im (**Fig 2I**). A large amount of CD8^+^ T cells were recruited into lung in dNS1-RBD group, and most of the infiltrated cells were virtual memory T cells (T_VM_) showing a CD44^+^CD62L^+^ phenotype with low CD49d expression (**Fig 2J-L**), which were recently reported to promote early viral control ^25^. The early infiltration of T_VM_ in the lung was significantly higher than that in NALT, lymph nodes, spleen, and other tissues (P < 0.05, **Fig 2M**). Together, these data suggest that dNS1-RBD and dNS1-Vector induce the activation and differentiation of innate immune cells (AMs, pDCs, NK cells and T_VM_) in lung tissue and form a local innate immune barrier.

### dNS1-RBD immunization reprogrammed chromatin accessibility of alveolar macrophages and maintained the trained immunity phenotype

In recent years, the memory effect of the innate immune response (trained immunity) is thought to play an important role in broad-spectrum anti-infection immunity ^26,27^. Trained AMs contribute to optimal protection against SARS-CoV-2 variants ^28^. High MHC II expression considered the trained phenotype of AMs ^29^. In this study, dNS1-RBD and dNS1-Vector significantly increased MHC II expression in mouse AMs, and no significant difference was found between the two groups **(Fig 3A)**. Furthermore, upregulation of CD80 and CD86 expression was observed in AMs indicating that AMs in the dNS1-RBD and dNS1-Vector groups were functionally changed as a result of innate immune training and maintaining an immunoreactive phenotype **(Fig 3B, C)**. ATAC-seq analysis was used to identify the potential changes that may occur in the chromatin accessibility of AMs induced by intranasal immunization. KEGG and GO enrichment analyses showed that differential ATAC-peaks were significantly enriched in pathways related to innate immune response and involved Toll-like- and retinoic acid inducible gene-1-like receptors (RLRs) (**Fig 3D**). A total of 202 upregulated and 194 downregulated peaks were detected in the dNS1-RBD group (**Fig 3E**). The dNS1-RBD group gained 810- and lost 760 lost open chromatin regions (OCRs), respectively compared with the control group 2 months after booster immunization in C57BL/6 mice (**Fig 3F**). Peaks that were not present in the control group were induced in both the dNS1-RBD and dNS1-Vector groups in the regulatory region of MHC II genes (*H2-Aaand H2-Eb1*) (**Fig 3G**). OCRs of several antiviral innate immune response genes including *TRIM25* and *IKBKB* are involved in RIG-I mediated antiviral signaling ^30,31^. Toll-like receptors, *TLR3* and *TLR1*, were not detected in the control group compared to the dNS1-RBD group (**Fig 3G**). AMs of dNS1-RBD hamsters gained and lost 913- and 1370 OCRs, respectively compared with the control at two weeks post prime vaccination (**Fig 3H**). The differential OCRs showed peaks in regulatory regions of glycolysis, pro-inflammatory cytokines, antiviral response, and toll-like receptor genes in both the dNS1-RBD and the dNS1-Vector group which were undetected in the control group (**Fig 3I**).

**Fig 3.**
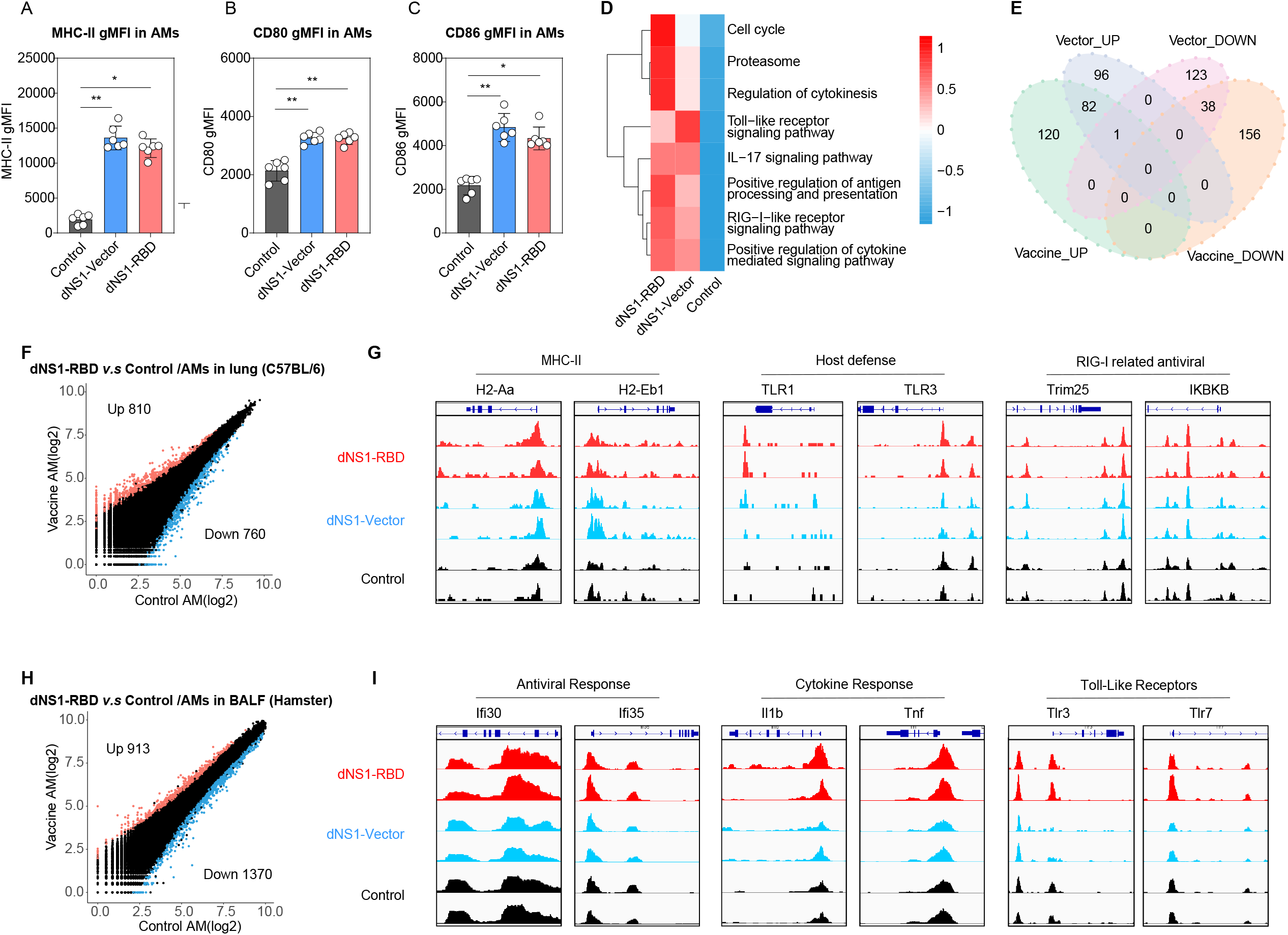
Trained phenotype of alveolar macrophages induced by dNS1-RBD and dNS1-Vector. (A-C) Statistical analysis plots for gMFI of MHC II (A), CD80 (B) and CD86 (C) on AMs in C57BL/6 mice. n = 6 mice/group. (D) KEGG and GO enrichment result of shared peaks, with color bar on –log_10_ (q-value) scale. (E) Venn-diagram showing differential ATAC-seq peaks (annotated as promoters) for dNS1-RBD or dNS1-Vector compared to the Control group. (F) Scatter-plot of differentially detected ATAC-seq peaks (log_2_FC > 1.5, q-value < 0.05) of AMs (Lung) in the dNS1-RBD vaccination group compared to the control group in C57BL/6. (G) IGV tracks showing differentially detected peaks related to host defense and antiviral response in C57BL/6 mice from dNS1-RBD vaccination group, dNS1-Vector vaccination group and control group. n = 2 mice/group. (H) Scatter-plot of differentially detected ATAC-seq peaks (log_2_FC > 1.5, q-value < 0.05) of AMs (BALF) in dNS1-RBD vaccination group compared to control group in hamsters. (I) IGV tracks showing differentially detected peaks related to host defense and antiviral response in hamsters. n = 2 mice/group. Data are presented as mean ± SEM. Statistical analysis for (A-C) were Kruskal-Wallis tests with Dunn’s multiple comparisons test. gMFI: Geometry Mean Fluorescence; AM: alveolar macrophages; Intensity, IGV: Integrative genomics viewer; BALF: bronchoalveolar lavage fluid.

These results suggest that NS1-impaired live attenuated influenza virus-induced trained immunity exerts nonspecific protection, possibly through epigenetic remodeling regulation and metabolic rewiring.

### dNS1-RBD immunization induced tissue-resident memory T cell responses covering the upper and lower respiratory tract

The adaptive immune response is established within days or weeks. Specific-T cells are indispensable for viral clearance and long-term immune protection. C57BL/6 mice were intranasally immunized with a single dose of dNS1-RBD or dNS1-Vector. Lung tissue mononuclear cells were harvested at 1-, 3-, 5- and 14 d.p.im to estimate the specific T cell response level. Intracellular cytokine (IFN-γ) expression was analyzed by flow cytometry after *ex vivo* stimulation using a 15-mer spike-peptide pool (**Fig S2A**). As expected, a specific T cell response was undetectable in the dNS1-Vector group (**Fig 4A-C**). In contrast, a CD8^+^ T cell response was detectable at 3 d.p.im in the dNS1-RBD group, with significant increasing numbers of IFN-γ^+^CD8^+^ T cells after 5 d.p.im (P < 0.01) (**Fig 4A-C**). The spleens were also analyzed to compare local- and systemic antigen-specific CD8^+^ T cell responses (**Fig S2B)**; the strength of the immune response (number and percentage of total immune cells) in spleen-specific T cells were weaker than that in lung tissues (**Fig S2C, D**). This correlated with enzyme-linked immunosorbent spot (ELISpot) assays of lymphocytes at 7 d.p.im after ex vivo antigen-peptide stimulation; NS1-RBD induced a robust T cell response in the lung tissue by producing 1974/SFC, which was 3.25 folds (606/SFC) that of the spleen, and 16.18 folds (122/SFC, P < 0.0001) that of the peripheral blood, respectively (Fig 4D, E). No specific T cell responses were generated on the first day in any tissue.

**Fig 4.**
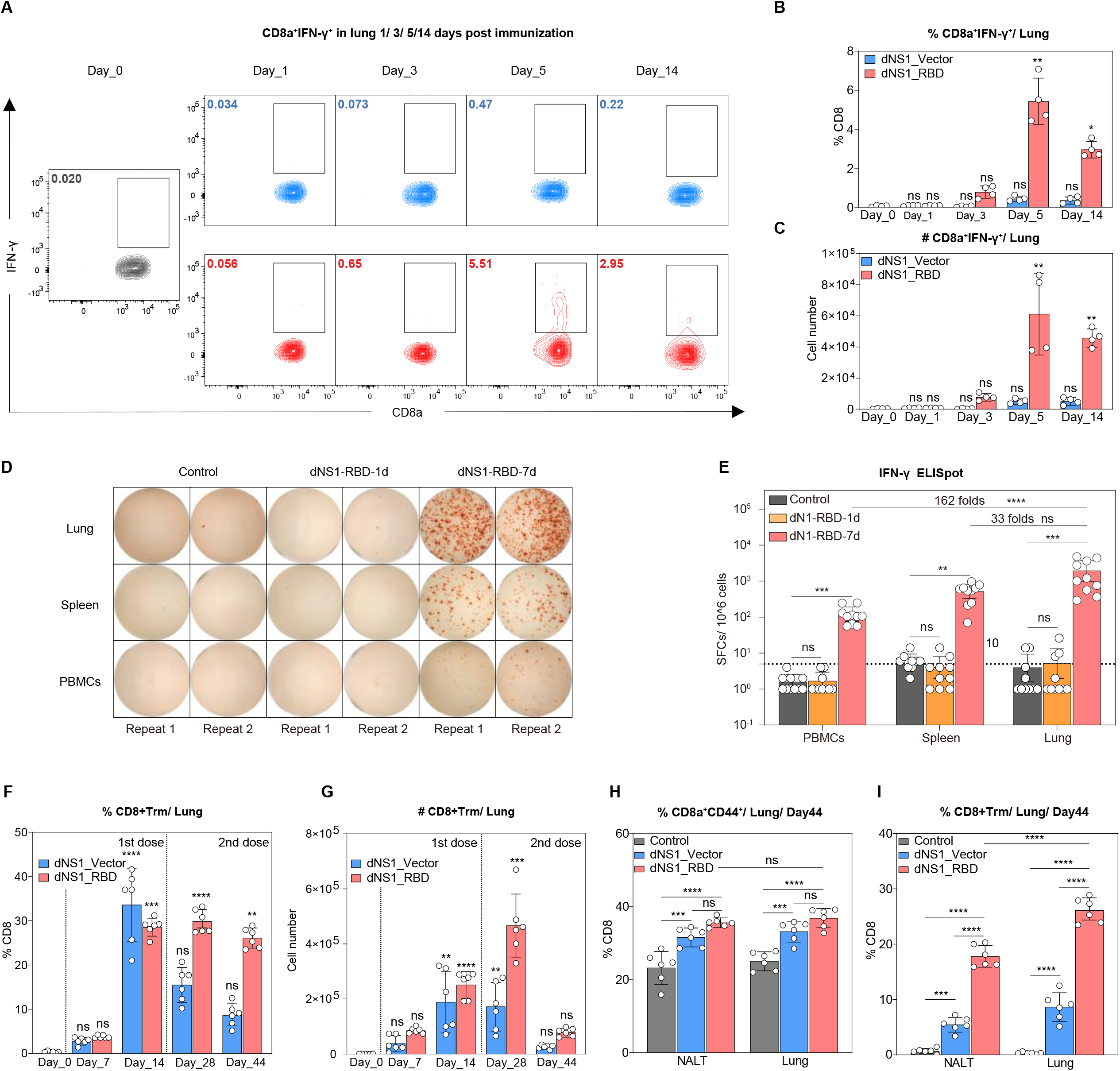
Intranasal immunization of dNS1-RBD and dNS1-Vector elicit tissue-resident memory T cells in Lungs. (A) Representative flow cytometry contour plots for CD8^+^ IFN-γ^+^ T cells staining. (B-C) Statistical analysis plots for percentage (B) and cell number (C) of CD8a^+^ IFN-γ^+^ in the lung. n = 4 mice/group. (D) Representative well images of the IFN-γ ELISpot response of the control group and dNS1-RBD group (1- and 7-days post-immunization). (E) The numbers of IFN-γ SFCs isolated from PBMCs, spleen, and lung were quantified after stimulation of a peptide pool covering the entire spike protein. n = 10 mice/group. (F-G) Bar graph depicting the frequency (F) and absolute number (G) of T_RMs_ in the lungs at indicated time points after prime and booster immunization. n = 6 mice/group. (H) Bar graph showing the frequency of CD44^+^ CD8^+^T cells in NALT and lung 30 days post a prime-boost vaccination. n = 6 mice/group. (I) Bar graph showing the frequency of CD8^+^ TRMs in NALT and lung 30 days post a prime-boost vaccination. n = 6 mice/group. Data are presented as mean ± SEM. Statistical analysis for (B, C, E, F, and G) were Kruskal-Wallis tests with Dunn’s multiple comparisons test. Statistical analysis for (H and I) were two-way ANOVA with Tukey’s multiple comparisons test. ns, non-significant. *p < 0.05, **p < 0.01, ***p < 0.001, ****p < 0.0001. IFN-γ: interferon-gamma; SFCs: spot-forming cells; PBMCs: peripheral blood mononuclear cells; T_RMs_: tissue-resident memory T cells; NALT: Nasal-associated lymphoid tissue.

Growing evidence supports a critical role for tissue-resident memory T cells (T_RMs_) in coordinating effective defense against reinfection in the local tissue where they reside ^32,33^. Single dose intranasal immunization using dNS1-RBD or dNS1-Vector CD8^+^ immediately activated T cells immediately and T_RMs_ were generated at 7 d.p.im **(Fig S2E**, and **S3A, B)**. Furthermore, potent induction of T_RMs_ in the lung tissue was observed by immunization with a second dose of dNS1-RBD at the subsequent time. However, boosting with the dNS1-Vector was less effective **(Fig 4F, G, and S3C, D)**. Similar observations were found with CD44^+^CD8^+^ T cells in the lungs. NALT and the lung tissue were harvested 30 days after the last immunization to verify the form of T_RMs_ in the upper respiratory tract. The frequency of CD44^+^CD8^+^ in the dNS1-Vector group was like that of the dNS1-RBD group. In contrast, dNS1-RBD induced greater levels T_RMs_ compared with the dNS1-Vector, particularly in the lung tissue (**Fig 2H, I** and **S3E, F)**.

Overall, intranasal immunization with dNS1-RBD induced RBD-specific T cell responses and tissue-resident memory T cell responses were concentrated in the respiratory tract.

### dNS1-RBD immunization provided protection from SARS-CoV-2 challenge in hamsters

Golden Syrian hamsters were challenged with beta SARS-CoV-2 variant by contact transmission to mimic patients with severe pneumonia caused by SARS-CoV-2 ^18^. Hamsters were sacrificed at 1-, 3-, 5 days post infection (dpi) for gross lung observation (**Fig 5A**). Control hamsters showed continuous body weight loss beginning at 1 dpi and exhibited weight loss of up to 11.97% at 5 dpi; in contrast, weight loss was not obvious in animals of dNS1-RBD group (mean: +1.24%) (**Fig 5B**). Vaccinated hamsters showed a lower viral RNA load in nasal washings at 1 dpi compared with the control hamsters. The significantly reduced viral RNA load (>2.0 log) in nasal washings, trachea and lung at 1 dpi (**Fig 5C**), suggesting that an early innate immune response may be elicited in the respiratory tracts and thus restrict SARS-CoV-2 replication. A relatively lower viral RNA load was observed at 3 dpi in trachea (> 1.0 log) and lung (> 2.0 log), which indicated that the intranasal vaccination inhibits the further infection of viruses from the upper to the lower respiratory tract of hamsters (**Fig 5C**). The lung viral loads in vaccinated hamsters were slightly lower than that of the control hamsters at 5 dpi although this was not statistically significant (**Fig 5C**). Hematoxylin and eosin (H&E) staining of the lung lobes of SARS-CoV-2-infected hamsters in the dNS1-RBD group showed significant alleviation of the pathological changes. In contrast, control animals exhibited typical features of severe pneumonia including increased lung lobe consolidation and alveolar destruction, diffusive inflammation, hyaline membrane formation, and severe pulmonary hemorrhage (**Fig 5D**). The apparent lesions in the dNS1-RBD group were markedly diminished, and no obvious viral-infection-related lung damage was observed in the gross lung images at 3- and 5 dpi compared with the control group (**Fig 5D**). The pathological severity scores of vaccinated hamsters were significantly lower than those of the control groups (**Fig 5C**). The reduced lung inflammation could have been linked with the immediate anti-viral responses of hosts ^34^.

**Fig 5.**
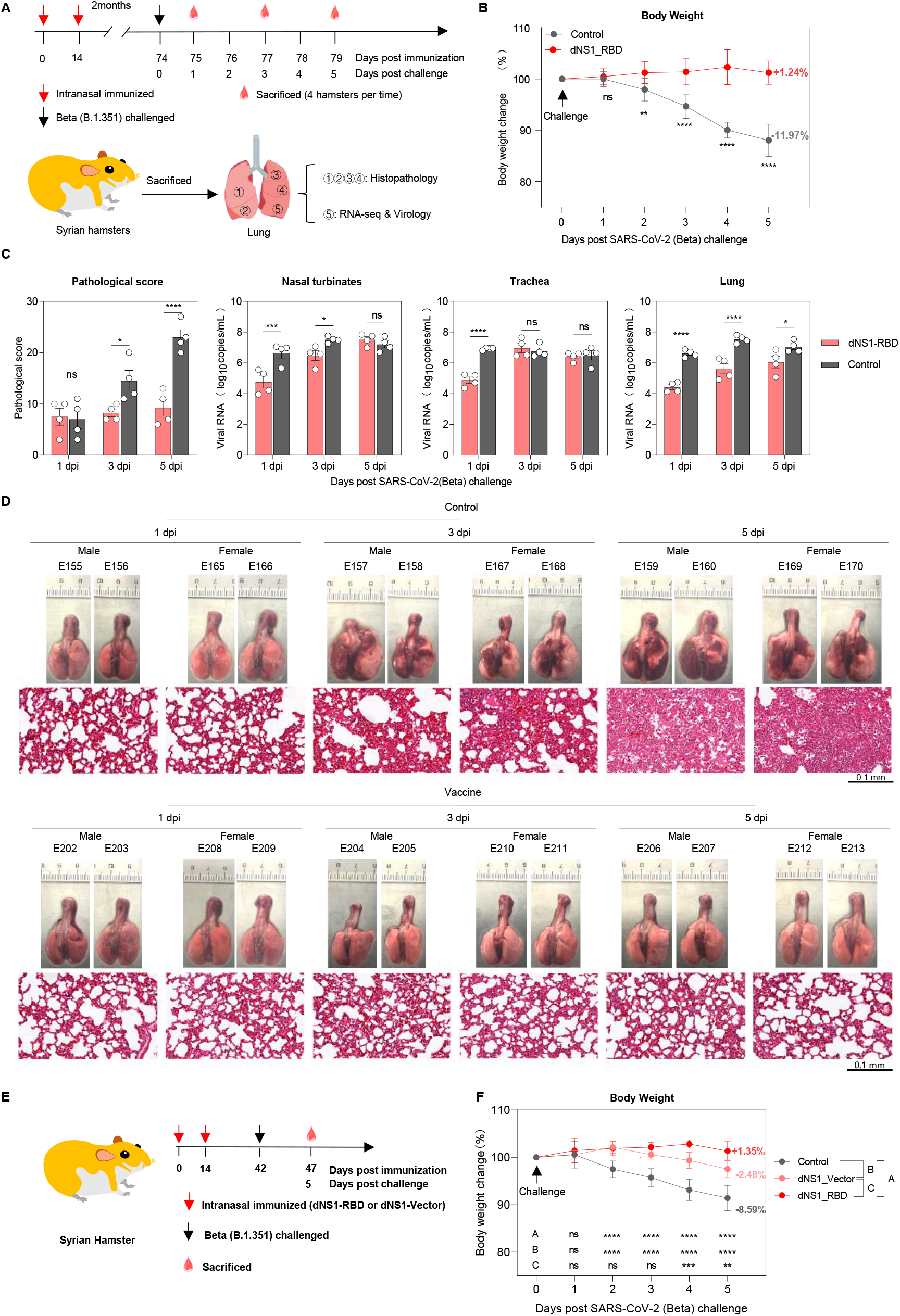
Intranasal immunization with dNS1-RBD and dNS1-Vector protect lung pathology in hamsters against SARS-CoV-2 Challenge. (A) Schema of the experimental design. On days 1, 3, and 5 after cohoused exposure, hamsters from vaccinated and control groups were euthanized for analyses. n = 4 hamsters/group. (B) Body weight changes of hamsters after cohoused exposure were plotted. The average weight loss of each group at 5 dpi is indicated as a colored number. (C) Bar graph showing the pathological severity scores of lungs and the viral RNA loads from nasal turbinate, trachea, and lung. (D) Gross lung images and H&E-stained lung sections from dNS1-RBD vaccinated and control groups. (E) Schema of the experimental design. Hamsters were intranasally vaccinated with dNS1-RBD or dNS1-Vector. (F) Body weight changes of hamsters after cohoused exposure were plotted. The average weight loss of each group at 5 dpi is indicated as a colored number. Data are presented as mean ± SEM. Statistical analysis for (B, C, and F) were two-way ANOVA with Bonferroni’s multiple comparisons test. n = 8 hamsters/group ns, non-significant. *p < 0.05, **p < 0.01, ***p < 0.001, ****p < 0.0001.

Hamsters were intranasally immunized with dNS1-RBD or dNS1-vector to validate whether a non-specific protective effect exist by infecting Beta variants through cohoused exposure (**Fig 5E**). The protective efficacy induced by dNS1-vector was comparable to that induced by the dNS1-RBD at early phase (1 and 2 dpi), but not later time point after infection (**Fig 5F**). Slightly enhanced protection was observed in hamsters vaccinated with dNS1-RBD, as reflected by their body weight change, with body weight change of +1.35% and −2.48% at 5 dpi respectively (**Fig 5F**). Altogether, hamsters from two vaccinated groups exhibited significantly improved weight loss when compared with the control hamsters. These data revealed that not only the dNS1-RBD but also the dNS1-vector confers protection independent of SARS-CoV-2-specific antibody and T-cell responses, suggesting that this NS1-deleted H1N1/CA4 vector-based intranasal vaccine could provide non-specific protection against respiratory virus. The cross-protection effects of vaccine could be achieved by trained immunity, with enhanced non-specific effector responses of innate immune cells.

### dNS1-RBD reduced inflammatory signaling- and pro-inflammatory factor levels post-challenge in hamsters

Distinct gene expression signatures visualized by PCA from hamster lung samples of the eight groups (pre-challenge and 1-, 3-, 5 dpi in the control group and the dNS1-RBD group) following SARS-CoV-2 infection showed tight clustering of biological samples. Control group samples were clearly separated in principal component 1 at 3 dpi and 5 dpi which contained 62.1% of the variation in the dataset. Meanwhile, all dNS1-RBD group samples and control group samples at 0- and 1 dpi were reflected in principal component 2 (6.8% of the variance). **(Fig 6A)**. Gene Ontology (GO) term analysis of the genes upregulated in control hamsters were enriched for response to virus, cytokine production, and inflammatory response regulation terms. Pro-inflammatory cytokines such as *IL-6, IL-1B* peaked at 3 dpi and remained high until 5 dpi in control hamster lungs **(Fig S4A)**. Excessive release of cytokines and overactivated antiviral responses may be associated with immunopathology upon infection. Notably, SARS-CoV-2 challenge did not significantly alter the cytokine profiles for dNS1-RBD vaccinated hamsters indicating that they were protected from over-activated inflammatory **(Fig S4B)**. DEGs involved in inflammatory cytokine production pathways such as *IL-6, IL-1B* and *IFN-*γ were elevated at 1 dpi, peaked at 3 dpi and remained high at 5 dpi in control hamsters based on KEGG pathway analysis (**Fig 6B)**. In contrast, the dynamics of transcription levels in the dNS1-RBD group changed to a much steadier state, with only mild elevation at 5 dpi (**Fig 6B)**. The heatmap showed that pathways such as the IFN response, TNF signaling and their downstream signaling pathway (for example, Jak-STAT and NF-κB signaling) were upregulated in control group yet remained steady in the dNS1-RBD group (**Fig 6B)**. Consistent with the lung pathology, Cell Death and Apoptosis pathway was clearly activated in control hamsters (**Fig 6C)**. Together, these cytokines and their related signaling pathways may play pathological roles in the initiation- and immune cell hyperactivation stage, ultimately leading to organ dysfunction in cytokine storms.

**Fig 6.**
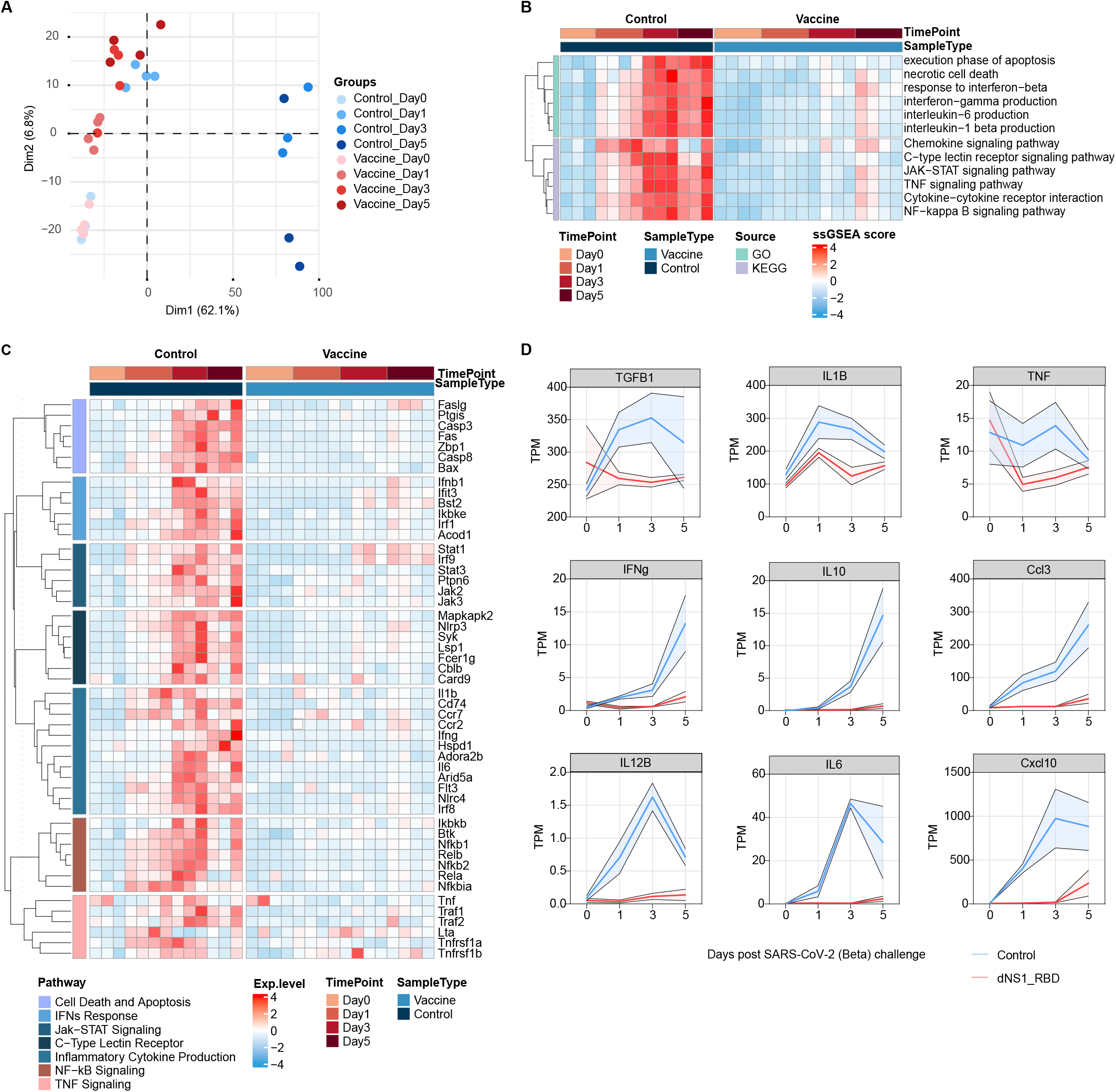
Transcriptome analysis of lung reveals distinct immune response between dNS1-RBD immunized and control hamster. (A) PCA Plot showing the global differences between vaccinated and control groups. (B) Heatmap visualization of scaled gene expression levels (TPM) for selected pathways of interest. (C) Heatmap showing the KEGG pathway enrichment analysis of DEGs. (D) Dynamic expression of cytokines in the lungs of hamsters, with error bars shaded (using standard error of the mean). PCA: Principal Component Analysis. n = 4 hamsters/group

Dysregulation of cytokines and chemokines is closely associated with severe inflammation, leading to tissue damage and destruction. After challenge, the expression of I*l-1b, Il-6, Cxcl10* rapidly increased and remained high until 5 dpi in the control group, which is consistent with the protein levels found in severe COVID-19 patients. Meanwhile, their levels remained at a relatively low level in the dNS1-RBD group (**Fig 6D**). Pro-inflammatory cytokines such as *IFN-*γ and *CCL3* showed a rapid and progressed elevation after SARS-CoV-2 exposure in control hamsters. Conversely, their expression levels remained at baseline levels in the dNS1-RBD group and did not progress in the later response despite the presence of the virus (**Fig 6D**). In addition, anti-inflammatory cytokines, such as *transforming growth factor-*β *(TGF-*β) and *IL-10* are two key immune homeostasis regulators which were relatively unperturbed after SARS-CoV-2 challenge (**Fig 6D**). Taken together, distinct responses to SARS-CoV-2 infection between the dNS1-RBD and control groups were identified in transcriptome signatures. Prior vaccination with dNS1-RBD alleviated the immunopathology caused by an over-activated inflammatory response. Overall, a finely tuned immune response prevents excessive inflammation and restores the homeostasis of the immune system and the organism.

## Discussion

Although intramuscular COVID-19 vaccines are widely used, controlling of the COVID-19 pandemic remains challenging. At the very beginning of the COVID-19 outbreak, our team began to develop an intranasal vaccine based on the NS1-deleted H1N1 vector carrying the gene encoding SARS-CoV-2-RBD (dNS1-RBD) ^18^. To our knowledge, this is the first intranasal spray COVID-19 vaccine that was entered into a phase 3 clinical trial ^19^ [ChiCTR2100051391]. In phases I and II clinical trials, the vaccine demonstrated good safety and a immunogenicity pattern that was highly consistent with animal studies—a weak peripheral immune response. Nevertheless, the vaccine is considered a seed candidate with a high probability of performing well in a phase III efficacy trial, based on the surprisingly strong, rapid, sustained and broad-spectrum protective results from animal studies, as well as the encouraging precedents of FluMist’s success ^35^. dNS1-RBD is characterized by the effective prevention of pathological changes caused by SARS-CoV-2 infection without inducing significant neutralizing antibodies ^18^, and dNS1-vector also showed protective effects (**Fig 5E, F**). This approach differs from the protective mechanism of traditional vaccines and is vastly different from the previous understanding of vaccine immunity.

The protective immune mechanism induced by the vaccine includes at least the following four aspects: (1) innate immunity (**Fig 1** and **Fig 2**), (2) trained immunity (**Fig 3**), (3) cellular immune responses covering the upper and lower respiratory tract (**Fig 4**), (4) antibody targeting RBD ^18^.

The innate immune system plays an important role in early infection control. The cellular signaling cascade can be activated through pattern recognition receptors of immune cells to promote cytokine- and chemokine secretion. Among these, the type-I interferon response is one of the first lines of defense against viral infections ^36^. After influenza A virus infection, its NS1 interacts with RIG-I and inhibits IFN-β production mediated by retinoic acid inducible gene-1(RIG-I), thereby allowing the virus to replicate in vivo. Therefore, NS1 truncation increases the host immune response, and includes the recruitment of innate immune cells and the production of various interferon-stimulated genes (ISGs) and cytokines, thereby enhancing early-stage immune protection ^37^. NS1 truncation promotes a stronger adaptive immune response ^38^. In our study, the innate immune response was observed in the lung tissue 24 h after dNS1-RBD vaccination through the activation of immune cells, multiple antivirals signaling pathways, and significant up-regulation of various cytokines including the RIG-I related pathway and a type-I interferon response (**Fig 1** and **Fig 2**). Our previous studies showed that the dNS1-RBD virus induced faster and stronger cytokine production than that of the wild-type influenza virus ^18^. Furthermore, there was a population of AMs called exudate lung macrophages with significantly upregulated CD11b, which we have not previously focused on. This is a common occurrence in post-infected lung tissue, wherein monocytes are recruited and rapidly differentiate into macrophages, which play an important role in remodeling alveoli and maintaining tissue homeostasis z. In addition, the virtual memory T cell (T_VM_, CD8^+^ CD44^+^ CD62L^+^ CD49d^low^) response participated in the early immune response to intranasal immunization (**Fig 2J, K**), which promotes early control of influenza virus infection ^25^. T_VM_ population also plays a role in bridging innate and acquired immune responses, which can differentiate into tissue resident-T cells. Rapid responses of these innate immune factors provide good protection within 24 h after dNS1-RBD vaccination; therefore, emergency vaccination in the early stage of the outbreak in the endemic areas may reduce disease burden by rapidly interrupting the spread of the virus.

RBD-specific cellular immune responses were induced in the NALT and the lung, and the number of IFN-γ spot-forming cells per million lymphocytes was approximately 16 times that in peripheral blood (**Fig 3D**). The response was generated on the 5^th^ day after vaccination and persisted for at least six months in the periphery ^18^. Earlier innate immune- and T cell responses are of great significance for asymptomatic infection or mild disease after SARS-COV-2 infection ^39–41^. Tissue-resident memory T cells (T_RMs_) in the respiratory tract and lungs are critical for controlling respiratory viral infections, and provide more timely-, and stronger protective immunity than circulating T cells ^32^. Lung tissue-resident T cells provide durable and broad-spectrum immune protection ^42–44^.

There is increasing evidence that innate immune cells can be modified by epigenetics to produce immune protection against heterologous pathogens within a certain period of time and exhibit trained immunity ^26,27,45^. dNS1-RBD vaccination can also alter the host response pattern to SARS-CoV-2 by inducing trained immunity with (1) broad-spectrum antiviral and (2) anti-inflammatory effects. It can relieve tissue inflammation and maintain tissue homeostasis, thereby providing pathological protection. dNS1-RBD activated alveolar macrophages, myeloid dendritic cells, and NK cells, and remodeled the chromatin openness of these cells. Genes related to anti-infection immunity in these cells remained open for several months, allowing for a faster and stronger response to SARS-CoV-2 challenge (**Fig 2** and **Fig 3)**. SARS-CoV-2 rapidly spread after challenge in the control group and reached 6.6 log_10_ copies/mL on the first day. This was accompanied by an over-aggressive immune response, and a variety of inflammation-related signaling pathways and factors were significantly upregulated. In contrast, hamsters in the dNS1-RBD group inhibited the copy number of SARS-CoV-2 virus by approximately 155-fold, 84-fold, and 10-fold on day1, 3, and 5, respectively (**Fig 5C**). Although the inhibitory effect was not prominent at day5, hamster lung tissue was healthy in the dNS1-RBD group, and transcriptome sequencing showed no overstimulated inflammatory response. Cytokines and chemokines are important for clearing viral infections; however, excessive inflammation leads to pathological damage. TGF-β, IL-6, IL10 are related to the occurrence of severe COVID-19 pneumonia. In addition, both viruses mainly infect upper respiratory tract epithelial cells and alveolar epithelial cells since the target cells infected by the dNS1-RBD virus highly overlap with those infected by SARS-CoV-2 ^46,47^, and widespread immune gene regulation in structural cells is reported ^48^. These studies suggest that dNS1-RBD may reshape the anti-infection response pattern by training immune cells and structural cells to attenuate inflammation and protect the organism.

Intranasal immunization with the NS1-shortened counterpart (A/PR8/NS124) provided good protection against lethal heterologous A/Aichi/2/68 (H3N2) influenza virus challenge, even without inhibition of the viral load, but significantly attenuated of inflammation and pathology ^16^. Interestingly, intranasal immunization with bacteria also showed some protection against the virus apart from the virus-against-virus effects. Intranasal immunization with *Autographa californica* nuclear polyhedrosis baculovirus (AcNPV) protected against lethal H1N1 A/PR/8/34 influenza virus challenge. Immune protection is observed 24 hours after immunization and the empty vector control also provide similar protection. This is very similar to our findings and indicates the importance of trained immunity in non-specific protection and its advantage in response speed ^49^. Intranasal immunization with attenuated *Bordetella pertussis* (BPZE1) protected against two different influenza A virus subtypes: H3N2 (A/Aichi/2/68) and H1N1 (A/ PR/8/34) through viral clearance, and alleviation of pathological symptoms by inhibiting the cytokine storm ^50^. Limited adaptive immunity in a BALB/c mouse model deficient in J segments (B cell KO) of the immunoglobulin heavy-chain locus with depleted CD4^+^ and CD8^+^ T cells (T cell dep) that was challenged with mouse-adapted virus (SARS-CoV-2 MA10) still showed immune protection suggesting that it is dependent on the trained immune function of alveolar macrophages ^15^.

Based on the findings of this study and previous knowledge, we suggest that the protective effects induced by intranasal vaccination could be not only dependent on viral clearance, but also through training the immune cells and structural cells in the respiratory tract, remodeling the immune microenvironment, and carrying out certain immunoregulatory effects after the heterologous challenge. This maintains the balance of the immune system and respiratory tissue and attenuates immune-induced tissue injury. Since intramuscular vaccines have been administered on a large worldwide, boosting with intranasal vaccines can establish more comprehensive immune protection in the anatomical space. At the same time, local cellular immunity and trained immunity in the respiratory tract are considered to be relatively broad-spectrum, which is beneficial for coping with the challenges caused by SARS-COV-2 variants.

## Materials and methods

### Animal experiments

All animal experiments strictly followed the recommendations of the Guide for the Care and Use of Laboratory Animals. The animal studies were approved by the Institutional Animal Care and Use Committee (IACUC) of Xiamen University.

The hamster studies were performed in an animal biosafety level 3 (ABSL-3) laboratory (State Key Laboratory of Emerging Infectious Diseases, The University of Hong Kong)

C57BL/6 mice were purchased from Shanghai SLAC Laboratory Animal Co.,Ltd. Golden Syrian hamsters were purchased from Beijing Vital River Laboratory Animal Technology Co., Ltd.

B6 (Jax 664) was originally from originally from Jackson Laboratory. All B6 (Jax 664) mice were maintained under SPF conditions before experiment, with experimental protocols approved by the Tsinghua Institutional Animal Care and Use Committee.

### Vaccine formulation

dNS1-RBD vaccine was prepared on a large scale at Beijing Wantai Biological Pharmacy Enterprise Co., Ltd., Beijing, China.

### Mouse immunization

Experimental animals were anesthetized with isoflurane, then intranasally immunized with 50 μL (1×10^6^ PFU/mL) of dNS1-RBD, whereas the control group was administered an equal volume of PBS. RNA-seq analysis involved intranasal immunization of C57BL/6 mice (three animals per group) with a single dose, and collection of lung tissue on day 1, 7, 14, or 28 after vaccination.

Innate immune response analyses involved intranasal immunization of C57BL/6 mice (4 animals per group) with a single dose, and collection of pulmonary lymphocytes on day 1, 3, and 5 after vaccination.

ELISpot analyses of peripheral blood mononuclear cells (PBMCs), splenic lymphocytes, pulmonary lymphocytes, and lymph node cells involved intranasal immunization of C57BL/6 mice with a single dose (10 animals per group), and pulmonary lymphocytes were collected on day 1 and 7 after vaccination.

Intracellular cytokine staining (ICS) analyses of pulmonary lymphocytes involved intranasal immunization of C57BL/6 mice with a single dose (four animals per group), and collection of pulmonary lymphocytes on day 1, 3, 5, and 14 after vaccination.

Tissue-resident T cell analyses involved intranasal immunization of C57BL/6 mice (6 animals per group) with a single dose and pulmonary lymphocytes were collected on day 7 and 14 after vaccination; another experiments schedule, intranasal immunization with a double dose (0/14 day) involved harvesting pulmonary lymphocytes on day 14 and 30 after boost vaccination.

### Hamster immunization and infection

The experimental hamsters (male: female = 1:1) were anesthetized with isoflurane and intranasally immunized with 100 μL of the vaccine (1×10^6^ PFU/mL), whereas the control group was administered with an equal volume of PBS. Two months after the last vaccination, hamsters were further evaluated by direct contact challenge with SARS-CoV-2. The donor hamsters (carrying the virus) were intranasally infected with 1× 10^3^ PFU of SARS-CoV-2. Each donor was transferred to a new cage and co-housed with the hamster of the dNS1-RBD group or control group for one day. The donor was then isolated, and the other hamsters were observed. Weight changes and typical symptoms (piloerection, hunched back, and abdominal respiration) were recorded daily after virus inoculation or contact. Hamsters were euthanized for tissue pathological and virological analyses and RNA-seq on day 1, 3, and 5 after challenge. Virus challenge studies were performed in an animal biosafety level 3 (ABSL-3) facility. The SARS-CoV-2 strain used in this study was B.1.351 variant AP100 (hCoV-19/China/AP100/2021; GISAID accession No. EPI_ISL_2779638)

### Organ-specific sample collection and organ dissociation

Mice were euthanized by exsanguination, and IACUC guidance was approved. Mice were transferred to a biosafety cabinet and their organs were carefully separated. All cells were counted using CountStar software.

### Lung

Lungs were cut into 0.5-cm pieces, placed in gentleMACS C tubes (Miltenyi) containing collagenase type IV (Gibco) and DNase I (Roche) in PBS containing 2% FBS, and dissociated using a gentleMACS Dissociator (Miltenyi; program m_lung_01). A single cell suspension was obtained by digesting tissue through a 70 µm cell strainer, and centrifugation at 300 g for 5 min at 4 °C. After centrifugation, 1 mL of cold red blood cell lysis buffer (Solarbio) was added for 2 min to lyse red blood cells. The reaction was stopped by adding 10 mL of cold PBS containing 2% FBS and washed once to remove residual red blood cell lysis buffer. Lymphocytes were obtained from the resulting cell suspensions using density gradient centrifugation (Percoll, SIGMA-ALDRICH). Cells were recovered at the interface of the 80% Percoll layer and the 40% Percoll layer, then washed with PBS + 2% BSA at 500 g for 5 min to remove excess Percoll.

### Lymph nodes

Cervical lymph nodes were carefully pinched with tweezers and rinsed several times with cold PBS containing 2% FBS. Lymph nodes were ground and passed through a 70 µm cell strainer. Lymphocytes were washed once and resuspended in PBS containing 2% FBS.

### Spleen

Mice were euthanized and their spleen were carefully separated and rinsed several times with cold PBS containing 2% FBS. Spleen were ground and passed through a 70 µm cell strainer and the cells were centrifuged at 300 g for 5 min at 4 °C. 10 ml of cold red blood cell lysis buffer (Solarbio) was added, and the samples were incubated for 5 min at 4 °C. The reaction was stopped by adding 20 ml of cold PBS containing 2% FBS and washed once to remove the residual buffer. Lymphocytes were washed once and resuspended in PBS containing 2% FBS.

### PBMCs

Mouse peripheral blood was transferred into a centrifuge tube containing sodium heparin, then 4 mL PBS buffer was added and transferred to SepMate™ PBMC isolation tubes (STEMCELL). PBMCs used density gradient centrifugation at 1200 g for 10 min at 25 °C (Ficoll-Paque PREMIUM, GE). PBMCs obtained from the middle layer cells.

### Nasal-associated lymphoid tissue (NALT)

The lower jaw of the mouse was removed, and a surgical knife was used to carefully cut and excise the upper palate by following the inner contour of mouse incisors and molar teeth. The tissue was digested at 37°C with collagenase type IV (Gibco) and DNase I (Roche) in PBS containing 2% FBS. A single-cell suspension was obtained from the digested tissue using a 70 µm cell strainer.

### Tissue-infiltrating lymphocytes

To distinguish tissue-infiltrating lymphocyte from cells associated with blood vessel, mice were intravenously injected with 1.5μg of fluorochrome conjugated anti-CD45.2 (clone 104) antibody 3 min before being sacrificed. The lung tissue was enzymatically digested in RPMI medium containing deoxyribonuclease I (20 μg/ml; Roche) and Liberase CI (40 μg/ml; Roche) at 37°C for 30 min. Lymphocyte was further isolated from tissue digest through Percoll density-gradient centrifugation (40/70%) at 1300g for 30 min. For flow cytometric analyses, lymphocytes prelabeled with intravenously injected antibodies were excluded by gating and unlabeled cells were termed ‘protected’.

### Flow cytometry

The expression of phenotypic markers, activation markers, and cytokines was evaluated. Briefly, cells were washed and blocked with antiCD16/CD32 (clone 2.4G2) in 0.5 % BSA-PBS for 30 min on ice, then stained with fluorochrome-labeled mAbs for 30 min on ice. ICS assays involved stimulating each sample with pooled spike peptides (1.0 μg/mL) in a U-bottom plate and incubating at 37 °C for 18 hours. GolgiPlug (BD Biosciences) was added to the culture at a final concentration of 1:1,000 and cells were further incubated for 6 additional hours. After incubation, cells were washed and stained with fluorochrome-labeled mAbs for 30 min on ice. The stained cells were fixed and permeabilized with BD Cytofix/Cytoperm (BD Biosciences, San Jose, CA, United States) according to the manufacturer’s instructions. The cells were washed and intracellularly stained with fluorochrome-labeled mAbs for 45 min on ice. The antibody reagents used in this study include: CD4 [Clone GK1.5, FTIC], CD8a [Clone 53-6.7, PE/Cy7], NK1.1 [Clone PK136, PerCP/Cy5.5], CD64 [Clone X54-5/7.1, PE /Cy7], CD170 [Clone S17007L, PE], CD11b [Clone M1/70, FITC], CD86 [Clone PO3, BV605], CD11c [Clone N418, BV421], CD45.2 [Clone 104, APC/Cy7], CD4 [Clone GK1.5, APC], CD8a [Clone 53-6.7, FITC], CD103 [Clone 2E7, PE], CD69 [Clone H1.2F3, BV421], CD44 [Clone IM7, PE/Cy7], CD45.2 [Clone 104, PerCP/Cy5.5], CD80 [Clone 16-10A1, PE /Cy7], CD11b [Clone M1/70, PE], CD317 [Clone 927,BV421], Ly-6C [Clone HK1.4, APC/Cy7], MHC class II [clone M5/114.15.2, APC], CD11c [Clone N418, APC]), cytokine expression (IFN-γ [clone XMG1.2, APC]), and a LIVE/DEAD® Fixable Aqua Dead cell stain kit was also used. Stained cells were processed using a BD LSRFORTESSA X-20 (BD Biosciences) flow cytometry system according to the manufacturer’s instructions. Data were analyzed using FlowJo X 10.0.7r2 and GraphPad Prism 8.

### ELISpot assay

Dissociated PBMCs, splenic lymphocytes, pulmonary lymphocytes, and lymph node cells were plated at 2.5×10^5^ into each well of a mouse IFN-γ ELISpot plate (Dakewe Biotech). Samples were stimulated using pooled Spike peptides of SARS-CoV-2 (Final concentration:1μg/mL, 15-mer peptide with 11 amino acids covering the spike region, Genscript) and cultured at 37°C with 5% CO_2_ for 20 h. Spots were scanned, counted, and quantified using the CTL S6 Universal Analyzer (Cellular Technology Limited) according to the manufacturer’s instructions.

### SARS-CoV-2 and dNS1-RBD RNA quantification

Detection of viral RNA levels was performed in hamster lungs using quantitative RT-PCR. Lung tissue was homogenized using TissueLyser II (Qiagen, Hilden, Germany) and RNA extraction was performed according to the manufacturer’s instructions (QIAamp Viral RNA mini kit, Qiagen). Viral RNA quantification was performed by measuring the copy number of the N gene using a SARS-CoV-2 RT-PCR kit (Wantai, Beijing, China), whereas CA4-dNS1-nCoV-RBD was quantified using RBD-targeted primers and the NS gene.

### Histopathology

Hamster lung tissues were fixed with 10% formalin for 48 h, embedded in paraffin, sectioned, and subjected to hematoxylin and eosin (H&E) staining. Whole-slide images of the lung sections were captured using a Leica Aperio Versa 200 microscope. Pathological lung lesions were scored based on i) Alveolar septum thickening and consolidation; ii) hemorrhage, exudation, pulmonary edema and mucous; iii) recruitment and infiltration of inflammatory immune cells. For each lobe, a score was determined based on the severity and percentage of injured areas. Four independent lobes of the lung tissues were scored and average lung pathological score of each individual hamster was used for pathological evaluation.

### Bulk RNA sequencing

The hamster lung lobe was removed, shredded into small pieces and stored in RNA Later Solution (Thermo Fisher Scientific) for a maximum of 24 h at 4 °C. Lung tissue was homogenized using TissueLyser II (Qiagen, Hilden, Germany) and RNA extraction was performed according to the manufacturer’s instructions (QIAamp Viral RNA Mini Kit (Qiagen)). The RNA samples were sent to OE Biotech Co., Ltd. (Shanghai, China) for RNA purification, cDNA library construction, and sequencing.

### ATAC sequencing

Hamster lungs were collected 2 months post-vaccination and sorted for bronchoalveolar lavage AMs using a BD FACS Aria Fusion machine. One hundred thousand sorted cells were centrifuged at 500 × g for 10 min at 4 °C per replicate. Cells were lysed with lysis buffer (10 mM Tris-HCl pH 7.4, 10 mM MgCl_2_, and 0.1% IGEPAL CA-630). Libraries were prepared using the TruePrep DNA library prep kit V2 for Illumina (Vazyme) according to the manufacturer’s instructions. Libraries were cleaned up with AMPure XP beads (Beckman coulter) at a ratio of 0.7 and the quality was assessed using the 2100 Bioanalyzer (Agilent Technologies). Libraries were sequenced with 150 paired ends using a NovaSeq 6000 instrument (Illumina) for an average of 20 million reads per sample.

### Analysis of bulk RNA-sequencing data

cDNA libraries were sequenced on an Illumina HiSeq X Ten platform and 150 bp paired-end reads were generated. Approximately 49.96 M raw reads were generated for each sample. Raw data (raw reads) in fastq format were initially processed using Trimmomatic and low-quality reads were removed to generate approximately 48.42 M clean reads for each sample for further analyses. The clean reads were mapped to the mouse genome (GRCm39) and hamster genome (BCM_Maur_2.0) using hisat2 version 2.2.1, then sorted using samtools version 1.15 for differentially expressed gene analysis. The raw count matrix was quantified using featureCounts version 2.0.1, and the transcripts per kilobase million (TPM) of each gene were calculated. Differential expression analysis was performed using R package DESeq2 version 1.34.0. A P value < 0.05, and foldchange > 2 was set as the threshold for significantly different expression. Hierarchical cluster analysis of differentially expressed genes (DEGs) was performed to determine the expression patterns in different groups and samples.

The enrichment analysis of DEGs through Gene Ontology (GO), Kyoto Encyclopedia of Genes and Genomes (KEGG) and gene-set enrichment analysis (GSEA) were performed using R package clusterprofile version 4.2.0 and fgsea version 1.20.0. Time-series analysis was performed using R package Mfuzz version 2.54.0. All visualizations related to RNA-seq analysis were made using R packages ggplot2 version 3.3.5, ComplexHeatmap version 2.10.0, and enrichplot version 1.14.1.

### Pre-processing and analysis of bulk ATAC-seq data

Quality control of the original ATAC-seq read file was performed using fasqc version 0.11.9 and multiqc version 1.12 software, and the raw data were trimmed using Trim_galore version 0.6.7 to remove the adaptors. The data were aligned to the GRCm39 and BCM_Maur_2.0 genome separately using Bowtie2 version 2.4.5 with the ‘--very-sensitive −X 2000’ parameter, followed by sorting using samtools version 1.15. Duplicated and unpaired reads were removed using the picard ‘MarkDuplicates’ command. Reads with mapping quality < 30, and reads aligned to the mitochondria chromosome were also removed. All downstream analyses were performed on the filtered reads. The bam file for all samples was converted to a bed file and then callpeak using MACS2 version 2.2.7.1 with the ‘-nomodel --shift -100 --extsize 200’ parameter.

Differential peak analysis was processed using bedtools to merge the peak file and featureCounts version 2.0.1 was used to construct the matrix; DESeq2 was then used to identify the differential peaks.

Coverage files from filtered bam files were produced using deeptools version 3.5.1 bamCoverage command. Each position was normalized with ‘—normalizeUsing RPGC,’ followed by conversion to bigWig format and visualization using IGV software.

## Acknowledgments

This study was supported by National Natural Science Foundation of China grants 81991491 (to N.X.), 32170943 (to T.Z.), 31730029 (to N.X.), and 82041038 (to Y.C.); Program on Key Research Project of China 2020YFC0842600 (to N.X.); Guangdong-Hongkong-Macau Joint Laboratory grant 2019B121205009 (to Y.G.); Natural Science Foundation of Fujian Province 2020J06007 (to T.Z.) and 2021J02006 (to Y.C.); Xiamen Youth Innovation Fund Project 3502Z20206060 (to T.Z.); and Fundamental Research Funds for the Central Universities 20720220003(to N.X.).

## Contributions

**Conceptualization**, J.Z., T.W., Y.C., T.Z., H.Q., Y.G., and N.X.; **Data curation**, L.Z., Y.J., J.H., J.C., R.Q., L.Y., and T.S.; **Formal analysis**, L.Z., Y.J., and J.H.; **Investigation**, L.Z., Y.J., J.H., J.C., R.Q., L.Y., T.S., X.L., Q.G., Y.C., C.Z., X.W., C.C., R.F., Y.W., F.C., H.X., and M.N. **Methodology**, L.Z., Y.J., J.H., J.C., R.Q., L.Y., T.S.; **Visualization**, L.Z., Y.J., J.H., J.C., R.Q., L.Y., and T.S.; **Validation**, L.Z., Y.J., and R.Q.; **Resources**, J.Z., Y.C., T.Z., H.Q., Y.G., and N.X.; **Funding acquisition**, Y.C., T.Z., Y.G., and N.X.; **Project administration**, T.Z.; **Supervision**, J.Z., T.W., Y.C., T.Z., H.Q., Y.G., and N.X.; **Writing-original draft**, L.Z., Y.J., J.H., and T.Z.; **Writing-review & editing**, J.Z., T.W., Y.C., T.Z., H.Q., Y.G., and N.X.

---

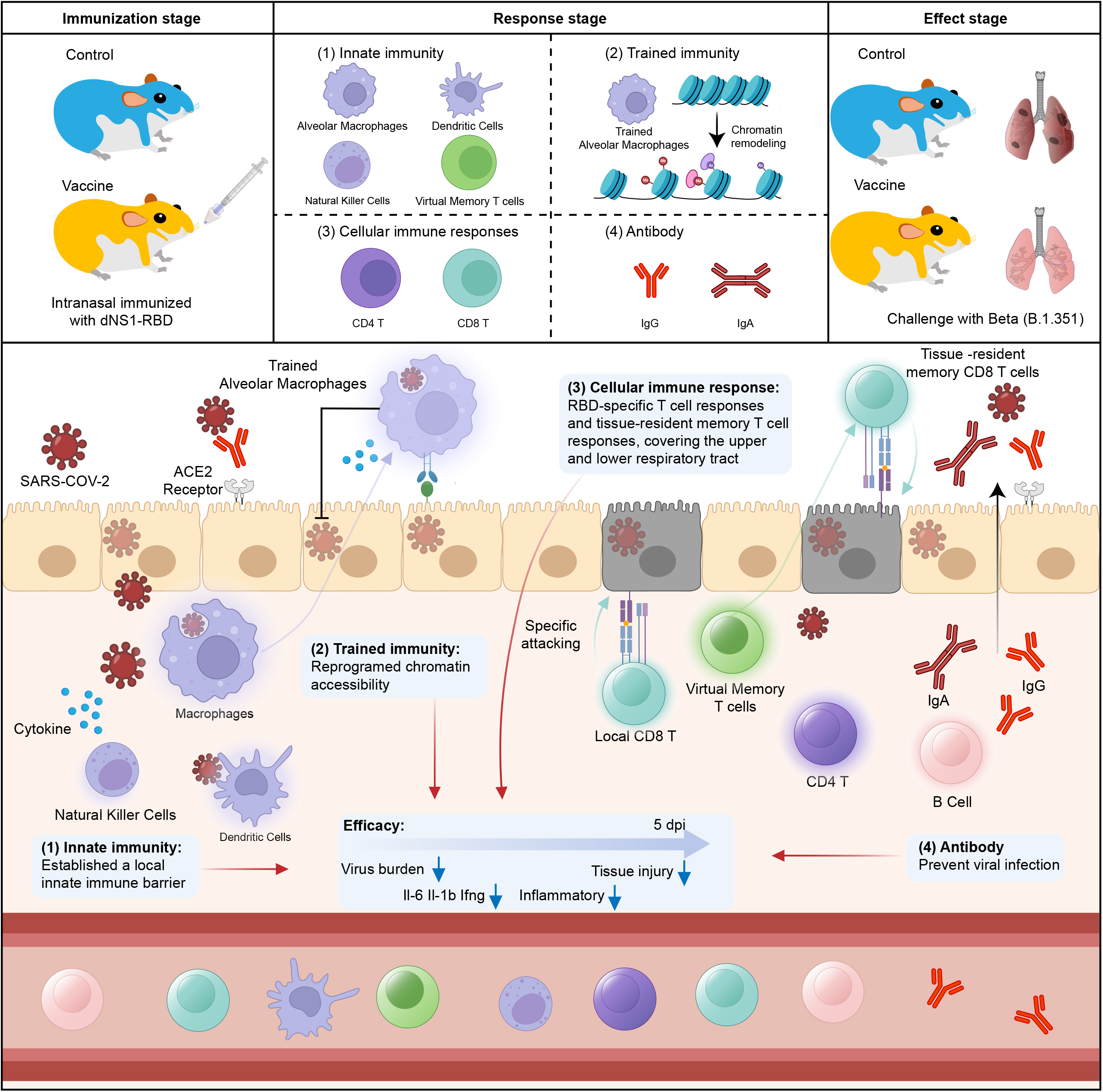

## Reference

1 Fontanet, A. & Cauchemez, S. COVID-19 herd immunity: where are we? Nat Rev Immunol 20, 583–584, doi:10.1038/s41577-020-00451-5 (2020).

2 Young, M., Crook, H., Scott, J. & Edison, P. Covid-19: virology, variants, and vaccines. BMJ Medicine 1, e000040, doi:10.1136/bmjmed-2021-000040 (2022).

3 Gruell, H. et al. Antibody-mediated neutralization of SARS-CoV-2. Immunity 55, 925–944, doi:10.1016/j.immuni.2022.05.005 (2022).

4 DeFrancesco, L. COVID-19 antibodies on trial. Nat Biotechnol 38, 1242–1252, doi:10.1038/s41587-020-0732-8 (2020).

5 Zhou, P. et al. A pneumonia outbreak associated with a new coronavirus of probable bat origin. Nature 579, 270–273, doi:10.1038/s41586-020-2012-7 (2020).

6 Sia, S. F. et al. Pathogenesis and transmission of SARS-CoV-2 in golden hamsters. Nature 583, 834–838, doi:10.1038/s41586-020-2342-5 (2020).

7 Harvey, W. T. et al. SARS-CoV-2 variants, spike mutations and immune escape. Nat Rev Microbiol 19, 409–424, doi:10.1038/s41579-021-00573-0 (2021).

8 Andrews, N. et al. Covid-19 Vaccine Effectiveness against the Omicron (B.1.1.529) Variant. N Engl J Med 386, 1532–1546, doi:10.1056/NEJMoa2119451 (2022).

9 Sokal, A. et al. Analysis of mRNA vaccination-elicited RBD-specific memory B cells reveals strong but incomplete immune escape of the SARS-CoV-2 Omicron variant. Immunity 55, 1096–1104.e1094, doi:10.1016/j.immuni.2022.04.002 (2022).

10 Shi, J. et al. Susceptibility of ferrets, cats, dogs, and other domesticated animals to SARS-coronavirus 2. Science 368, 1016–1020, doi:10.1126/science.abb7015 (2020).

11 Chandler, J. C. et al. SARS-CoV-2 exposure in wild white-tailed deer (Odocoileus virginianus). Proc Natl Acad Sci U S A 118, doi:10.1073/pnas.2114828118 (2021).

12 Alu, A. et al. Intranasal COVID-19 vaccines: From bench to bed. EBioMedicine 76, 103841, doi:10.1016/j.ebiom.2022.103841 (2022).

13 Hassan, A. O. et al. A Single-Dose Intranasal ChAd Vaccine Protects Upper and Lower Respiratory Tracts against SARS-CoV-2. Cell 183, 169–184.e113, doi:10.1016/j.cell.2020.08.026 (2020).

14 Mettelman, R. C., Allen, E. K. & Thomas, P. G. Mucosal immune responses to infection and vaccination in the respiratory tract. Immunity 55, 749–780, doi:10.1016/j.immuni.2022.04.013 (2022).

15 Afkhami, S. et al. Respiratory mucosal delivery of next-generation COVID-19 vaccine provides robust protection against both ancestral and variant strains of SARS-CoV-2. Cell 185, 896–915.e819, doi:10.1016/j.cell.2022.02.005 (2022).

16 Vasilyev, K., Shurygina, A. P., Sergeeva, M., Stukova, M. & Egorov, A. Intranasal Immunization with the Influenza A Virus Encoding Truncated NS1 Protein Protects Mice from Heterologous Challenge by Restraining the Inflammatory Response in the Lungs. Microorganisms 9, doi:10.3390/microorganisms9040690 (2021).

17 Wang, P. et al. Generation of DelNS1 Influenza Viruses: a Strategy for Optimizing Live Attenuated Influenza Vaccines. mBio 10, doi:10.1128/mBio.02180-19 (2019).

18 Chen, J. et al. A live attenuated virus-based intranasal COVID-19 vaccine provides rapid, prolonged, and broad protection against SARS-CoV-2. Sci Bull (Beijing), doi:10.1016/j.scib.2022.05.018 (2022).

19 Zhu, F. et al. Safety and immunogenicity of a live-attenuated influenza virus vector-based intranasal SARS-CoV-2 vaccine in adults: randomised, double-blind, placebo-controlled, phase 1 and 2 trials. Lancet Respir Med, doi:10.1016/s2213-2600(22)00131-x (2022).

20 Grant, R. A. et al. Circuits between infected macrophages and T cells in SARS-CoV-2 pneumonia. Nature 590, 635–641, doi:10.1038/s41586-020-03148-w (2021).

21 Califano, D., Furuya, Y. & Metzger, D. W. Effects of Influenza on Alveolar Macrophage Viability Are Dependent on Mouse Genetic Strain. J Immunol 201, 134–144, doi:10.4049/jimmunol.1701406 (2018).

22 Milanez-Almeida, P. et al. CD11b(+)Ly6C(++)Ly6G(-) Cells with Suppressive Activity Towards T Cells Accumulate in Lungs of Influenza a Virus-Infected Mice. Eur J Microbiol Immunol (Bp) 5, 246–255, doi:10.1556/1886.2015.00038 (2015).

23 Kim, J. et al. Low-dielectric-constant polyimide aerogel composite films with low water uptake. Polymer Journal 48, 829–834, doi:10.1038/pj.2016.37 (2016).

24 Lanzer, K. G., Cookenham, T., Reiley, W. W. & Blackman, M. A. Virtual memory cells make a major contribution to the response of aged influenza-naïve mice to influenza virus infection. Immun Ageing 15, 17, doi:10.1186/s12979-018-0122-y (2018).

25 Hou, S. et al. Virtual memory T cells orchestrate extralymphoid responses conducive to resident memory. Sci Immunol 6, eabg9433, doi:10.1126/sciimmunol.abg9433 (2021).

26 Netea, M. G. et al. Trained Immunity: a Tool for Reducing Susceptibility to and the Severity of SARS-CoV-2 Infection. Cell 181, 969–977, doi:10.1016/j.cell.2020.04.042 (2020).

27 Mantovani, A. & Netea, M. G. Trained Innate Immunity, Epigenetics, and Covid-19. N Engl J Med 383, 1078–1080, doi:10.1056/NEJMcibr2011679 (2020).

28 Xing, Z. et al. Innate immune memory of tissue-resident macrophages and trained innate immunity: Re-vamping vaccine concept and strategies. J Leukoc Biol 108, 825–834, doi:10.1002/jlb.4mr0220-446r (2020).

29 Yao, Y. et al. Induction of Autonomous Memory Alveolar Macrophages Requires T Cell Help and Is Critical to Trained Immunity. Cell 175, 1634–1650.e1617, doi:10.1016/j.cell.2018.09.042 (2018).

30 Timmer, A. M. & Nizet, V. IKKbeta/NF-kappaB and the miscreant macrophage. J Exp Med 205, 1255–1259, doi:10.1084/jem.20081056 (2008).

31 Liu, Z. et al. NDR2 promotes the antiviral immune response via facilitating TRIM25-mediated RIG-I activation in macrophages. Sci Adv 5, eaav0163, doi:10.1126/sciadv.aav0163 (2019).

32 Zens, K. D., Chen, J. K. & Farber, D. L. Vaccine-generated lung tissue-resident memory T cells provide heterosubtypic protection to influenza infection. JCI Insight 1, doi:10.1172/jci.insight.85832 (2016).

33 Szabo, P. A., Miron, M. & Farber, D. L. Location, location, location: Tissue resident memory T cells in mice and humans. Sci Immunol 4, doi:10.1126/sciimmunol.aas9673 (2019).

34 Plaçais, L. et al. Immune interventions in COVID-19: a matter of time? Mucosal Immunol 15, 198–210, doi:10.1038/s41385-021-00464-w (2022).

35 Treanor, J. J. et al. Evaluation of trivalent, live, cold-adapted (CAIV-T) and inactivated (TIV) influenza vaccines in prevention of virus infection and illness following challenge of adults with wild-type influenza A (H1N1), A (H3N2), and B viruses. Vaccine 18, 899–906, doi:10.1016/s0264-410x(99)00334-5 (1999).

36 Hoagland, D. A. et al. Leveraging the antiviral type I interferon system as a first line of defense against SARS-CoV-2 pathogenicity. Immunity 54, 557–570.e555, doi:10.1016/j.immuni.2021.01.017 (2021).

37 Vasilyev, K., Yukhneva, M., Shurygina, A., Stukova, M. & Egorov, A. Enhancement of the immunogenicity of influenza A virus by the inhibition of immunosuppressive function of NS1 protein. Microbiology Independent Research journal 5, 48–58, doi:10.18527/2500-2236-2018-5-1-48-58 (2018).

38 Vasilyev, K. A., Shurygina, A.-P. S., Stukova, M. A. & Egorov, A. Y. Enhanced CD8+ T-cell response in mice immunized with NS1-truncated influenza virus. Microbiology Independent Research journal 7, 24–33, doi:10.18527/2500-2236-2020-7-1-24-33 (2020).

39 Tan, A. T. et al. Early induction of functional SARS-CoV-2-specific T cells associates with rapid viral clearance and mild disease in COVID-19 patients. Cell Rep 34, 108728, doi:10.1016/j.celrep.2021.108728 (2021).

40 Blanco-Melo, D. et al. Imbalanced Host Response to SARS-CoV-2 Drives Development of COVID-19. Cell 181, 1036–1045.e1039, doi:10.1016/j.cell.2020.04.026 (2020).

41 Bergamaschi, L. et al. Longitudinal analysis reveals that delayed bystander CD8+ T cell activation and early immune pathology distinguish severe COVID-19 from mild disease. Immunity 54, 1257–1275.e1258, doi:10.1016/j.immuni.2021.05.010 (2021).

42 Wu, T. et al. Lung-resident memory CD8 T cells (TRM) are indispensable for optimal cross-protection against pulmonary virus infection. J Leukoc Biol 95, 215–224, doi:10.1189/jlb.0313180 (2014).

43 Turner, D. L. et al. Lung niches for the generation and maintenance of tissue-resident memory T cells. Mucosal Immunol 7, 501–510, doi:10.1038/mi.2013.67 (2014).

44 Teijaro, J. R. et al. Cutting edge: Tissue-retentive lung memory CD4 T cells mediate optimal protection to respiratory virus infection. The Journal of Immunology 187, 5510–5514, doi:10.4049/jimmunol.1102243 (2011).

45 Netea, M. G. et al. Trained immunity: A program of innate immune memory in health and disease. Science 352, aaf1098, doi:10.1126/science.aaf1098 (2016).

46 Flerlage, T., Boyd, D. F., Meliopoulos, V., Thomas, P. G. & Schultz-Cherry, S. Influenza virus and SARS-CoV-2: pathogenesis and host responses in the respiratory tract. Nat Rev Microbiol 19, 425–441, doi:10.1038/s41579-021-00542-7 (2021).

47 Sungnak, W. et al. SARS-CoV-2 entry factors are highly expressed in nasal epithelial cells together with innate immune genes. Nature Medicine 26, 681–687, doi:10.1038/s41591-020-0868-6 (2020).

48 Thomas Krausgruber, N. F., Victoria Fife-Gernedl, Martin Senekowitsch, Linda C Schuster, Alexander Lercher, Amelie Nemc, Christian Schmidl, André F Rendeiro, Andreas Bergthaler, Christoph Bock. Structural cells are key regulators of organ-specific immune responses. Nature 583, 296–302, doi:10.1038/s41586-020-2424-4 (2020).

49 Abe, T. et al. Baculovirus induces an innate immune response and confers protection from lethal influenza virus infection in mice. J Immunol 171, 1133–1139, doi:10.4049/jimmunol.171.3.1133 (2003).

50 Li, R. et al. Attenuated Bordetella pertussis protects against highly pathogenic influenza A viruses by dampening the cytokine storm. J Virol 84, 7105–7113, doi:10.1128/jvi.02542-09 (2010).

