## Supplemental for "Intranasal delivery of NS1-deleted influenza virus vectored COVID-19 vaccine restrains the SARS-CoV-2 inflammatory response"

**Supplementary Materials**

**Fig. S1**

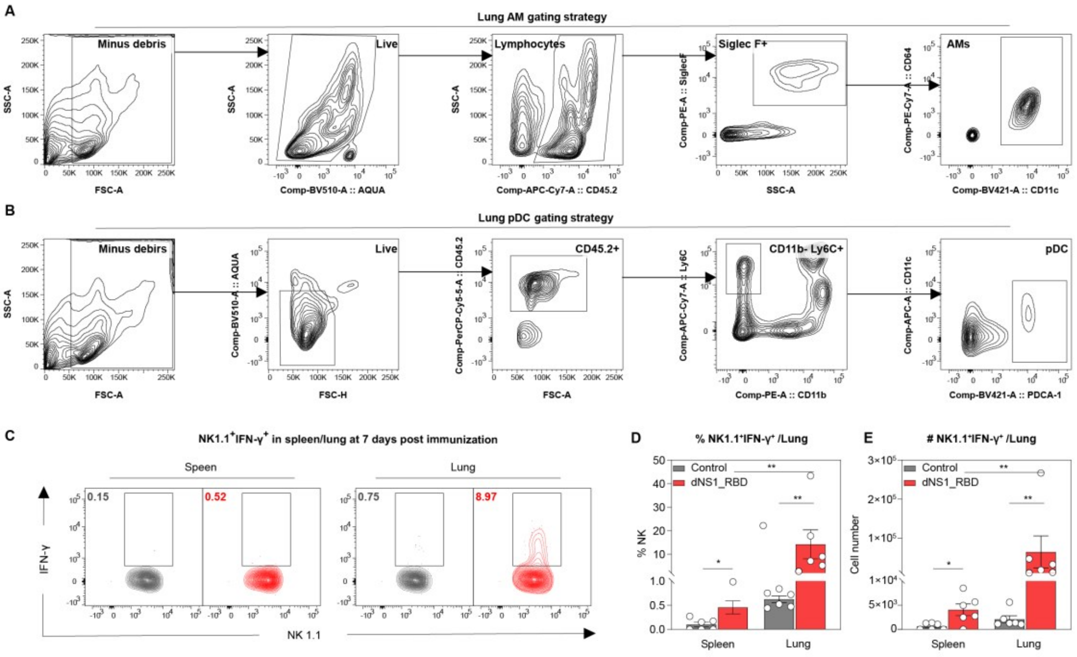


**Fig. S1 dNS1-RBD induces production of interferon-gamma in NK1.1+ cells.**

(A) Gating strategy for AMs in the lung.

(B) Gating strategy for pDCs and NK cells in the lung.

(C) Flow cytometry analysis of IFN-γ-producing NK1.1$^+$-cells in the lung and spleen at 7 days post-immunization.

(D-E) Bar graph depicting frequency (D) and absolute number (E) of NK1.1$^+$ IFN-γ$^+$ cells in the lung and spleen at 7 days post-immunization. n = 6 mice/group.

Data are presented as mean ± SEM. Statistics were Mann-Whitney tests. ns, non-significant. *$p < 0.05$, **$p < 0.01$, ***$p < 0.001$, ****$p < 0.0001$.

AMs: alveolar macrophages; pDCs: plasmacytoid dendritic cells; NK cells: natural killer cells; IFN-γ: interferon-gamma.

**Fig. S2**

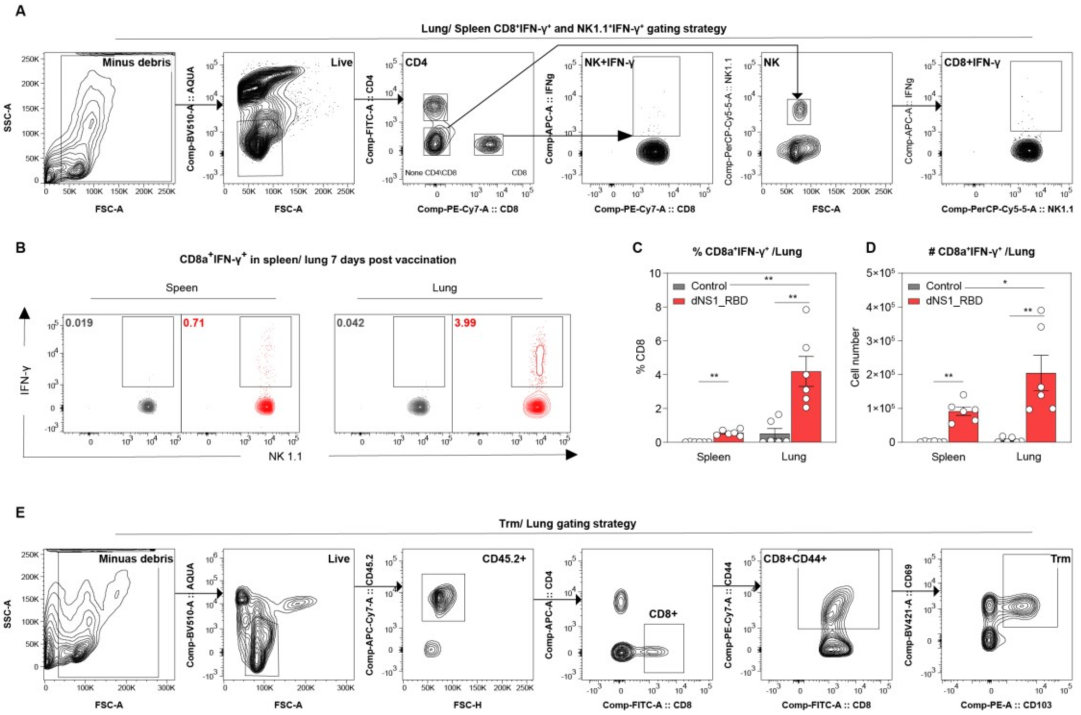


**Fig. S2 dNS1-RBD induces SARS-CoV2-specific T-cell response in mice.**

(A) Gating strategy for NK cells and IFN-γ$^+$ CD8$^+$ T cells in the lung and spleen.

(B) Representative flow cytometry contour plots showing IFN-γ produced by CD8$^+$ T cells in the lung and spleen at 7 days post-immunization.

(C-D) Bar graph showing frequency (D) and absolute number (E) of IFN-γ-producing CD8$^+$ T cells in the lung and spleen. n = 6 mice/group.

(E) Gating strategy for CD44$^+$ CD8$^+$ T cells and TRMs in the lung and NALT.

Data are presented as mean ± SEM. Statistics analysis were Mann-Whitney tests.

ns, non- significant. *p < 0.05, **p < 0.01.

NK cells: natural killer cells; IFN-γ: interferon-gamma; $T_{RMs}$: tissue-resident memory T cells; NALT: Nasal-associated lymphoid tissue.

30 **Fig. S3**

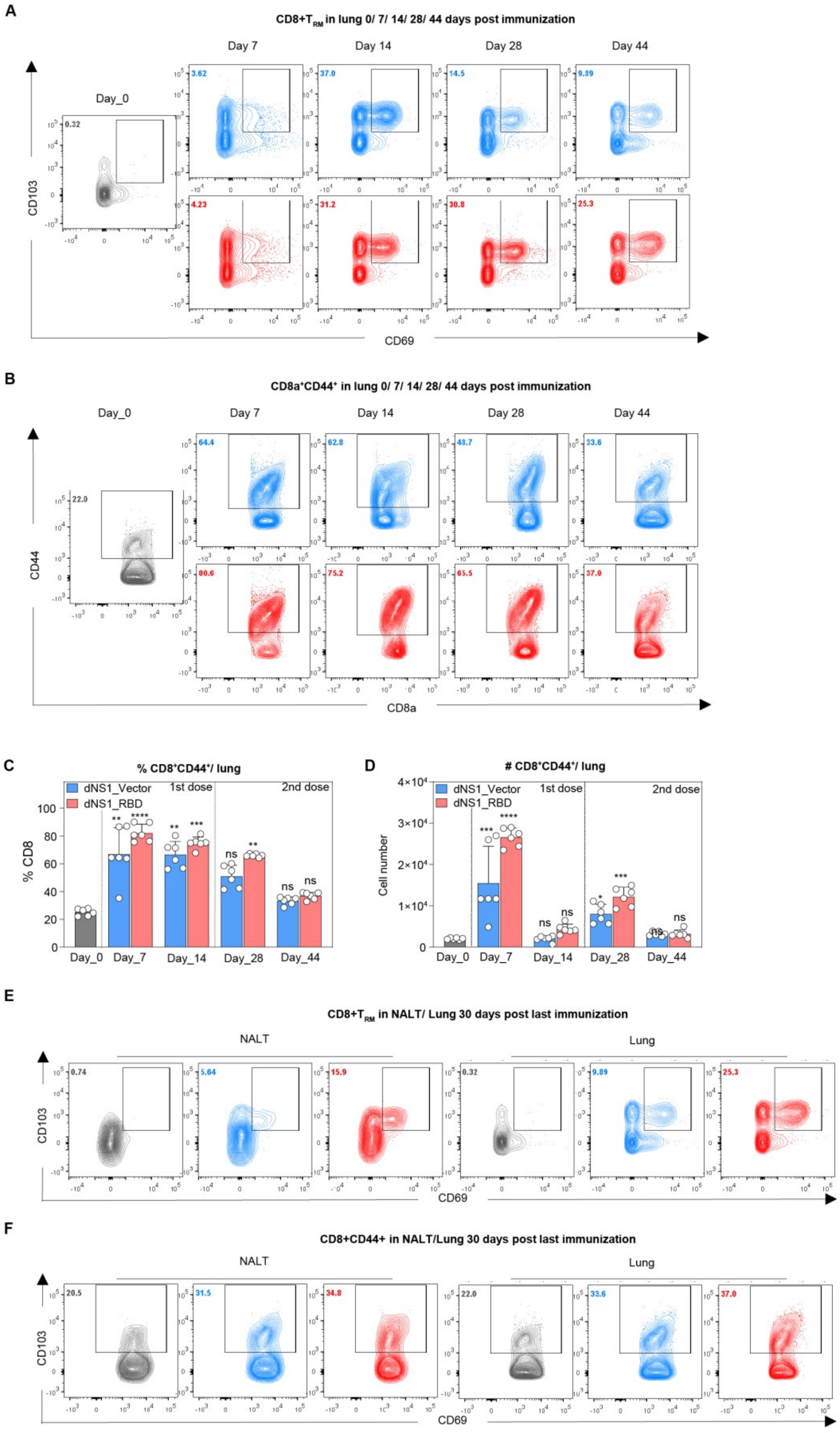

A
CD8+T_RM in lung 0/ 7/ 14/ 28/ 44 days post immunization
Day_0
Day 7
Day 14
Day 28
Day 44
CD103
CD69
B
CD8a+CD44+ in lung 0/ 7/ 14/ 28/ 44 days post immunization
CD44
CD8a
C
% CD8+CD44+/ lung
dNS1_Vector
dNS1_RBD
1st dose
2nd dose
% CD8
Day_0
Day_7
Day_14
Day_28
Day_44
D
### CD8+CD44+/ lung
Cell number
E
CD8+T_RM in NALT/ Lung 30 days post last immunization
NALT
Lung
CD103
CD69
F
CD8+CD44+ in NALT/Lung 30 days post last immunization
NALT
Lung
CD103
CD69


31

**Fig. S3 dNS1-RBD induces tissue-resident memory T cells in mice.**

(A) Representative flow cytometry contour plots for $T_{RMs}$ in the lung.

(B) Representative flow cytometry contour plots for $CD44^+$ $CD8^+$ T cells in the lung.

(C-D) Bar graph showing frequency (C) and absolute number (D) of $CD44^+$ $CD8^+$ T cells in the lung at indicated time points. n = 6 mice/group.

(E) Representative flow cytometry contour plots for $CD8^+$ $T_{RMs}$ in NALT and lung.

(F) Representative flow cytometry contour plots for $CD44^+$ $CD8^+$T cells in NALT and lung.

Data are presented as mean ± SEM. Statistical analysis for (C and D) were Kruskal-Wallis tests with Dunn's multiple comparisons test.

ns, non- significant. *p < 0.05, **p < 0.01, ***p < 0.001, ****p < 0.0001.

$T_{RMs}$: tissue-resident memory T cells; NALT: Nasal-associated lymphoid tissue.

**Fig. S4**

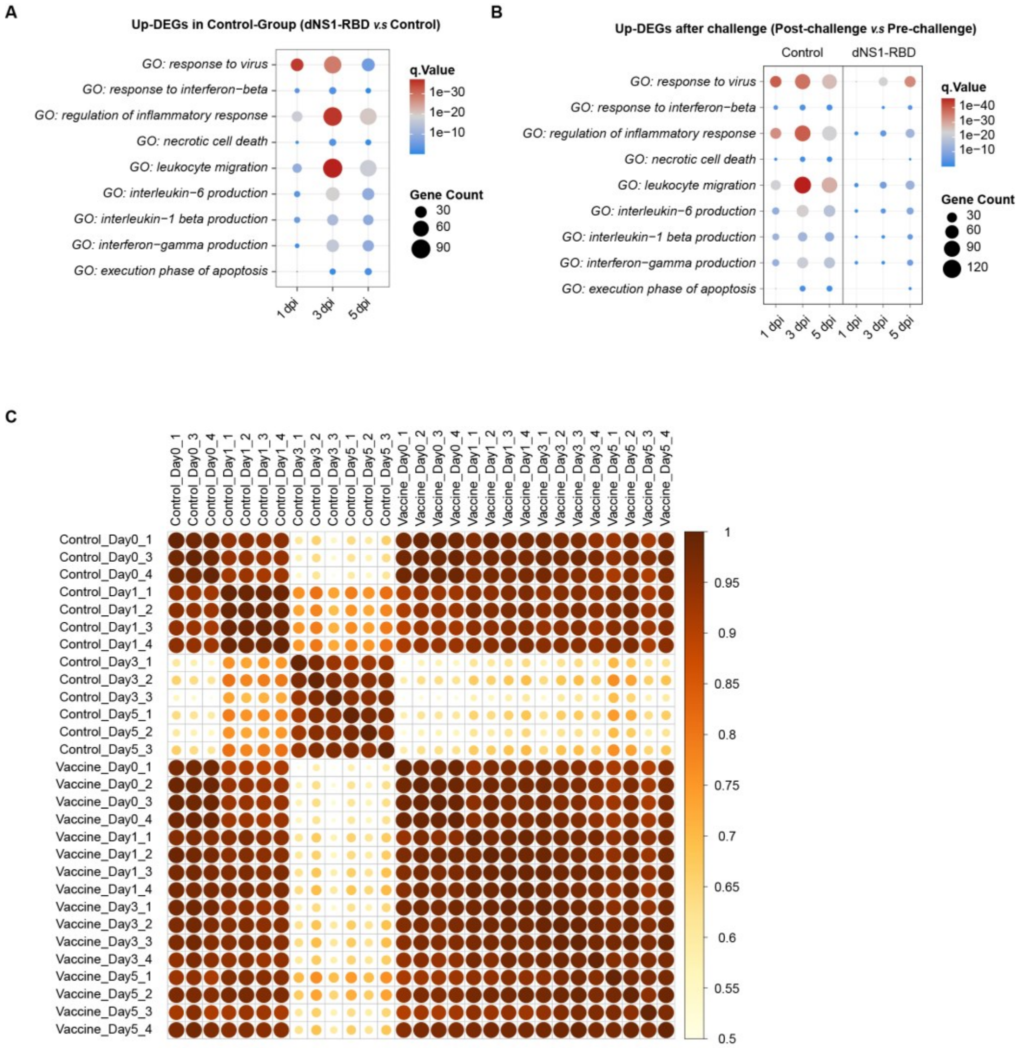


**Fig. S4 Intranasal vaccination with dNS1-RBD attenuated the inflammation responses after SARS-Cov-2 challenge.**

(A) Dot plots showing the GO enrichment analysis of up-regulated differentially expressed genes (Up-DEGs) related to immune responses in control hamsters compared to vaccinated hamsters.

(B) Dot plots showing the GO enrichment analysis of up-regulated differentially expressed genes (Up-DEGs) related to immune responses compared to 0 dpi in control hamsters and vaccinated hamsters respectively.

(C) Pearson correlation analysis between all biological RNA-sample.
